# Searching α−solenoid proteins involved in organellar gene expression

**DOI:** 10.1101/2025.09.09.674916

**Authors:** Céline Cattelin, Rebecca Goulancourt, Emanuel Chatellet, Alexis Astatourian, Charles Robert, Francis-André Wollman, Ingrid Lafontaine

**Affiliations:** UMR7141, Photobiology and Physiology of Plastids and Microalgae, P3M, Sorbonne Université, CNRS, Institut de Biologie Physico-Chimique, 13 rue Pierre et Marie Curie, F-75005 Paris, France; UPR9080, Laboratoire de Biochimie Théorique, Université Paris Cité, CNRS, Institut de Biologie Physico-Chimique, 13 rue Pierre et Marie Curie, 75005 PARIS, France

## Abstract

In photosynthetic eukaryotes of the green lineage, the expression of the chloroplast genome is mainly regulated post-transcriptionally, by RNA-binding proteins encoded in the nuclear genome (OTAF for Organelle Trans-Acting Factor). Most of those identified to date belong to two families of α−solenoid proteins - the pentatrico-peptide repeat (PPR) and octatrico-peptide repeat (OPR) families - and interact with specific sequences on their target mRNAs through a domain composed of repeated motifs, allowing their maturation, splicing, editing, stabilization and translation activation. To identify new OTAFs, we developed three approaches for annotating α- solenoid proteins targeted to the chloroplast or the mitochondria. One to identify distant homologs of existing OTAF families, and two others (decision tree and random forest classifiers) to identify new OTAF families. The combined approaches efficiently retrieve previously annotated OTAFs in 2 *model organisms.* It identified 1067 OPR proteins and 4983 PPR proteins in 43 proteomes of Archaeplastida. Our analysis also identified putative proteins composed of both OPR and PPR domains. Finally, our results identified 3300 other α-solenoid candidates which are likely to participate as new regulators of organelle gene expression. In particular, we identified new candidates in species in which the regulatory mechanisms of plastid gene expression are still understudied, such as in the glaucophyte *Cyanophora paradoxa* and the red alga *Porphyridium purpureum*. We thus provide valuable new tools to decipher the repertoire of OTAF, as well as new candidates for experimental characterization in the entire eukaryotic tree of life.

## Introduction

Eukaryotic photosynthesis is ensured by plastids, organelles originally acquired *ca.* 1.5 billion years ago from a primary endosymbiosis involving a protist host and a cyanobacterial ancestor, which gave rise to the extant green algae and land plants (together Viridiplantae), to rhodophyte and glaucophyte algae (Archibald 2015). These endosymbiotic events were followed by massive gene transfers from the plastid progenitors to the nucleus of the host cell, as it had already happened during mitochondrial endosymbiosis, which probably involved an Archaeal host and an α-proteobacterial ancestor, *ca.* 1.8 billion years ago (Archibald 2015). Most of the organelle proteome - either mitochondrion or plastid-, is now nucleus-encoded, translated in the cytosol and imported in the organelle. However, these energy-providing organelles have retained a tiny genome, therefore most of the major protein complexes associated with their bioenergetic membranes are genetic mosaics with subunits encoded in two different genomes. To ensure cell viability and acclimation of the organelle activity (energy production and metabolism), expression of organelle genomes became, during evolution, closely interconnected with that of the host cell. While the expression of nuclear genes is regulated at multiple levels (transcriptionally and post-transcriptionally as well as via epigenetic marks), plastidial and mitochondrial genomes are known to date to be mainly regulated post-transcriptionally by nucleus-encoded RNA-binding proteins, as demonstrated by pioneering studies in Viridiplantae and in yeast (Rochaix 1992; Fox 1996; Choquet and Wollman 2002). Most of the factors described up to now, hereafter named OTAF (for organellar trans-acting factor), contain a succession of either PPR motifs of 35 residues (pentatrico-peptide repeat) or OPR motifs of 38 residues (octatrico-peptide repeat) (Small and Peeters 2000; Eberhard et al. 2011; Barkan and Small 2014; Hammani et al. 2014). They belong to a large class of proteins containing repeated motifs forming anti-parallel α-helix pairs, like HEAT, Ankyrin, Armadillo and Pumilio repeats, which confer a rod-shape like structure with concave surface where ligands can bind (Andrade and Bork 1995; Riggleman et al. 1989; Spassov and Jurecic 2003; Mosavi et al. 2004). PPR and OPR motifs are degenerated and form pairs of antiparallel α-helices. The variable number of repeats of the motif form an α-solenoid shape with a positively charged surface that binds to the mRNA in a sequence-specific manner (Barkan and Small 2014; Cheng et al. 2016). PPR repeats are related to TPR repeats (Sikorski et al. 1990; Small and Peeters 2000) that mainly mediate protein-protein interactions and are involved in a variety of cell processes (D’Andrea and Regan 2003). Sel1-like repeats are themselves linked to TPR repeats, while HAT repeats are “Half-A-TPR” repeats, some of which, like PPR, bind to RNA (Preker and Keller 1998). The PPR and OPR protein repertoires are remarkably diverse between organisms, with land plants containing several hundreds of PPR and few OPR, whereas the green alga *Chlamydomonas reinhardtii* encodes only a dozen PPR (Tourasse et al. 2013; Gutmann et al. 2020) but more than a hundred of OPR (Eberhard et al. 2011; Rahire et al. 2012; Merendino et al. 2006; Marx et al. 2015; Lefebvre-Legendre et al. 2015; Wang et al. 2015). PPR and OPR proteins also contain additional domains, like the DYW domain in PLS-type PPR, involved in editing (Manna 2015; Barkan and Small 2014; Hayes and Santibanez 2020) and the RAP domain with a probable endonuclease activity (Boehm et al. 2017) at the C terminus of several OPR in *C. reinhardtii* (Eberhard et al. 2011; Boulouis et al. 2015; Lefebvre-Legendre et al. 2015).

To tackle the issue of the PPR motifs degeneracy, profile-based methods have been developed, like TPRpred that identifies TPR motifs and related PPR and Sel1-like motifs (Karpenahalli et al. 2007); PPRFinder based on plant homologs (Gutmann et al. 2020) and SCIPHER based on yeast homologs (Lipinski et al. 2011). These approaches use taxonomically restricted profiles of either plants or yeasts to perform motif search. Given the highly biased distribution of PPR and OPR repeats across the eukaryotic tree, it is likely that those families are still incomplete, especially for OPR.

Here, we present three procedures to complete the catalogue of known OTAFs and discover new OTAF families, which we applied to a representative set of Archaeplastida species. The first approach is an iterative profile-based similarity procedure to retrieve distant OTAFs homologs **p**redicted to be **t**argeted to **o**rganelles (*pto*), in order to fully describe their distribution. Applied to OPR and PPR, we show that OPR expansions were restricted within Chlorophyta and that outside of Viridiplantae, PPRs and OPRs are few in number, suggesting that other players in the regulation of gene expression in organelles remain to be discovered. These differences likely reflect genetic adaptation to different lifestyles or ecological niches. We also present two machine learning procedures to retrieve new families of nuclear-encoded candidates likely to be involved in organelle genome expression, *i.e. pto* proteins adopting an α-solenoid shape with similar physico-chemical properties as known OTAFs. We thus identified several dozens of new *pto* α-solenoid candidates, including a family of α-solenoid proteins tandemly duplicated in *C. reinhardtii*, whose experimental characterization would be relevant to understand their possible contribution to chloroplast gene expression.

The tools developed in this study will allow the identification of OTAF outside Archaeplastida, which is of genuine interest, owing to the scarce knowledge on the regulation of gene expression in organelles beyond what has been reported in Opistokhonta and Viridiplantae.

## Results

### Procedures for the identification of OTAF candidates

We developed procedures to identify distant homologs of known OTAF families, and to identify novel OTAF candidates. These procedures were applied to establish the OTAF catalogues within a set of 43 proteomes from all three taxa of Archaeplastida: the only available one in Glaucophyta, the 5 available ones in Rhodophyta, and within Chloroplastida, the 23 Chlorophyta reference proteomes available at Uniprot, plus the JGI proteome of *C. reinhardtii*, and 12 proteomes of Streptophyta including *A. thaliana* (Table S1).

#### Iterative Profile-based Procedure to retrieve homologs with known motifs

To retrieve distant homologs of known OTAFs, we developed an Iterative Profile-Based Procedure (IPB) (Figure 1, left panel). Profiles built from a defined set of motifs (for example PPR or OPR motifs) are searched against the proteomes of interest to identify new motifs. Proteins with significant hits against one or more profile are selected if they are predicted to be localized in mitochondria or chloroplast (pto) by at least 2 over 4 prediction algorithms (see Methods). The procedure can be iterated a defined number of times, or until no new *pto* protein is identified. After each iteration, all motifs (newly identified and already existing) are clustered according to their similarity into clustersmotifs with Markov Clustering (see Methods). A new profile is built for each clustermotif, and all obtained profiles will be used for the next iteration. In this study, we built profiles from 107 motifs from 12 published OPR proteins and 155 P-motifs, the most abundant type of PPR motifs (Cheng et al. 2016) from 11 PPR proteins (Table S2). With the OPR profiles, 10 iterative steps were performed to reach convergence, *i.e.* until no new *pto* protein was retrieved. With the PPR motifs, 10 iteration steps nearly reached convergence (only one protein was retrieved after the last iteration).

**Figure 1:**
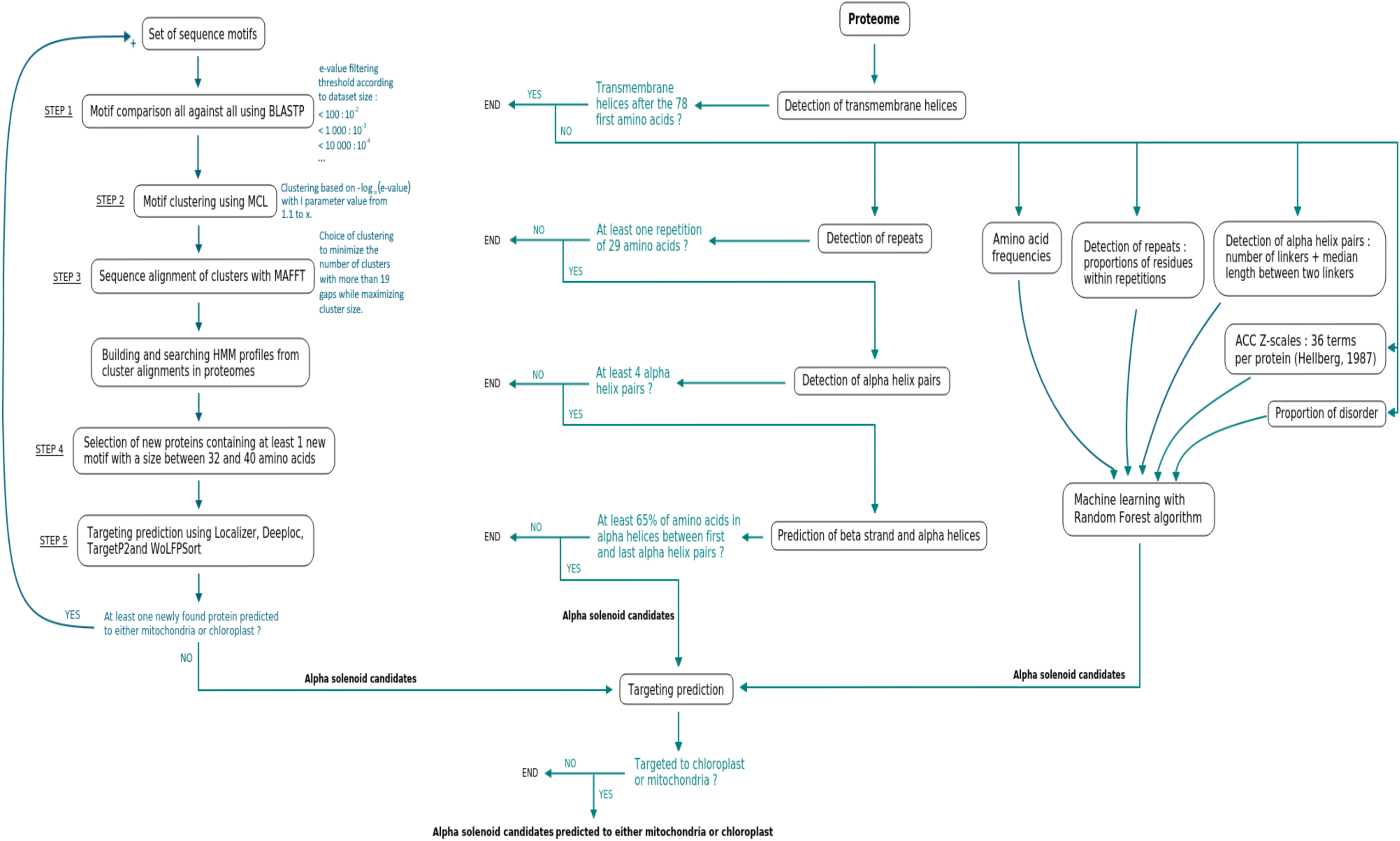
Workflow of the three developed procedures. IPB, left; DT, middle; RF, right

In order to find new OTAF candidates with physico-chemical and structural properties similar to those of the known OTAF but not necessarily evolutionary related to OPR and PPR families, we developed two classification procedures based on predicted protein properties, one based on a decision tree and one based on a random forest. These approaches thus retrieve potential α- solenoid proteins composed of repeated motifs and pairs of α-helices targeted to an endosymbiotic organelle.

#### Decision tree procedure

The decision-tree procedure (DT) is based on both the detection of repeated sequences (sequence repeat) within the protein and of predicted anti-parallel α-helix pairs (structural repeat), because the sequence similarity between the structural repeats could be undetectable and because a sequence repeat does not necessarily fold into an anti-parallel α-helix pair (Methods and Figure 1, central panel). All proteins containing a predicted transmembrane helix by TMHMM (see Methods) are excluded, as most of the published works describe OTAFs as soluble proteins. Only candidates with at least two repeated sequences (>=26 amino acids), *i.e.* two motifs predicted by RADAR, were selected. Selected candidates at the third step are those that also contain at least 4 linkers (separated by 32 up to 400 amino acids) predicted by ARD2 (Fournier et al. 2013). A linker is defined as the amino acid(s) between the two α-helices of a structural repeat unit. Four linkers implies that there are four structural repeats, not necessarily detected as sequence repeats, or three structural repeats if ARD2 detects a linker between structural units. The maximum distance of 400 amino acids between two linkers implicitly takes into account the fact that ARD2 can miss some linkers (Fournier et al. 2013). In that perspective, 400 amino acids would correspond broadly to the maximum distance between the last and the first repeat of a suite of 10 structural repeats.

The parameterization of DT was performed on a training set composed of 426 α-solenoid proteins (positive control) and 286 non α-solenoid proteins (negative control) classified based on their experimentally resolved or predicted 3D structure (see Methods). Threshold values for RADAR and ARD2 have been determined to maximize the precision and recall of the detection on the training set (Table 1), but the F1-score of DT remains modest (0.46), with a rather good specificity 0.73 at the expense of recall (0.34) (Table 1).

**Table 1.**
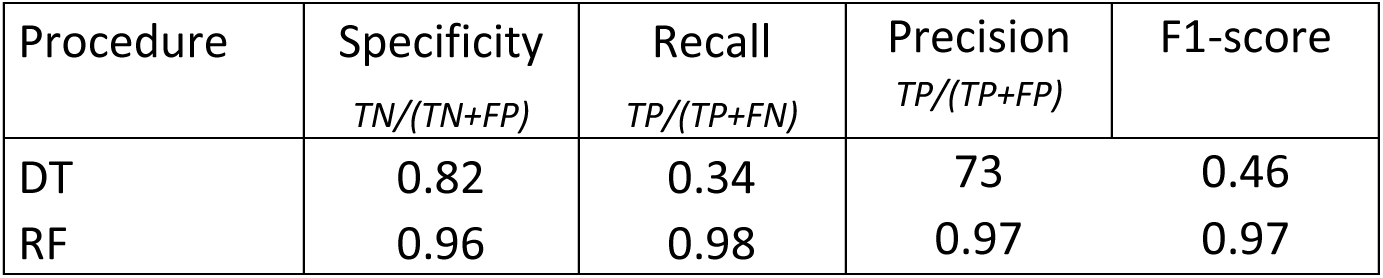
Performances of IPB and DT on the training set and average performance of 1000.

| Procedure | Specificity<br>*TN/(TN+FP)* | Recall<br>*TP/(TP+FN)* | Precision<br>*TP/(TP+FP)* | F1-score |
| --- | --- | --- | --- | --- |
| DT | 0.82 | 0.34 | 73 | 0.46 |
| RF | 0.96 | 0.98 | 0.97 | 0.97 |

#### Random forest procedure

To improve the sensitivity of the similarity-free detection approach and to avoid the definition of threshold values to select candidates like in the DT approach, we also developed a random forest (RF) classifier (Figure 1, right panel), in which composition and physico-chemical properties are also considered. Each protein is thus described by 61 variables. Four variables directly describe α−solenoid properties: the proportion of amino acids in repeats, the median length of the repeats, the number of linkers between two α-helices and the median length between the predicted linkers. The remaining general variables are the 20 amino-acid frequencies proportion of amino acids with disorder propensity, and 36 Auto-Cross Correlation terms between the Z-scales physico-chemical descriptors for each amino-acid (Hellberg et al. 1987) as described in (Garrido et al. 2020). The RF classifier was trained on the same training set as the DT procedure (see Methods). Over 1000 iterations of the model on our validation test (those 10% proteins from the positive and negative sets that were never used to train the model), on average the precision is 0.97, sensitivity 0.98 and specificity 0.96 (Table 1). For detection, RF was run 1000 times and candidates were retained if they were retrieved in at least 90% of the iterations.

The importance of each variable in the classifier is shown on Figure S1. The most important one is the proportion of residues within repeat regions, and the second ones are the AAC term z1.2.lag4, that reflects amphipathic constraints and the frequency of Proline, which is known to destabilize the conformation when occurring in the middle of an α-helix, but with a stabilizing effect at their N-terminus (Kim and Kang 1999). The median length of the repeats plays only a limited role. This was quite expected given the wide distributions in both positive and negative controls (see Methods). The importance of variables related to the α-helices (number of linkers and median length between linkers) and the proportion of disorder is lower than 32/60 other variables, making RF notably different from DT.

Comparative ability of the three procedures to retrieve those OPR and PPR proteins, previously annotated in model organisms.

We determined how many of the annotated PPR in *A. thaliana* and OPR in *C. reinhardtii* were retrieved by each procedure.

#### Iterative Profile-Based

From P-class PPR motifs, IPB retrieved 95% (451/475) of the PPR proteins listed in the PPR database at University of Western Australia (Cheng et al. 2016) with a UniProtKB identifier in the *A. thaliana* proteome (named PPR UWA hereafter) and 3 PPR not listed at UWA, but annotated as PPR in Swissprot. IPB also retrieved 10 of the 14 PPR annotated in *C. reinhardtii* (Table 2). Regarding OPR proteins, IPB retrieved the unique one (ATRAP, AT2G31890) in *A. thaliana* and 119 already annotated OPR in *C. reinhardtii*. Among those OPR in *C. reinhardtii*, 56 (out of 58) are published OPR proteins and 57 (out of 60) additional proteins annotated as “OctotricoPeptide Repeat Protein” in the annotation v5.6 version available at JGI, Phytozome13 (Table 2). There are 5 OPR that IPB failed to retrieve, most probably because their OPR motifs are too distant from the ones with used: the published OPR Raa3 (Rivier et al. 2001) and NCL18 (Boulouis et al. 2015) and three additional OPR annotated at JGI (Cre07.g347500 annotated as OPR114, Cre02.g146900 and Cre07.g336500). However, they were retrieved by RF and or DT (see below). In summary, the IPB procedure has 95% precision and 93% sensitivity (Table 3).

**Table 2.**
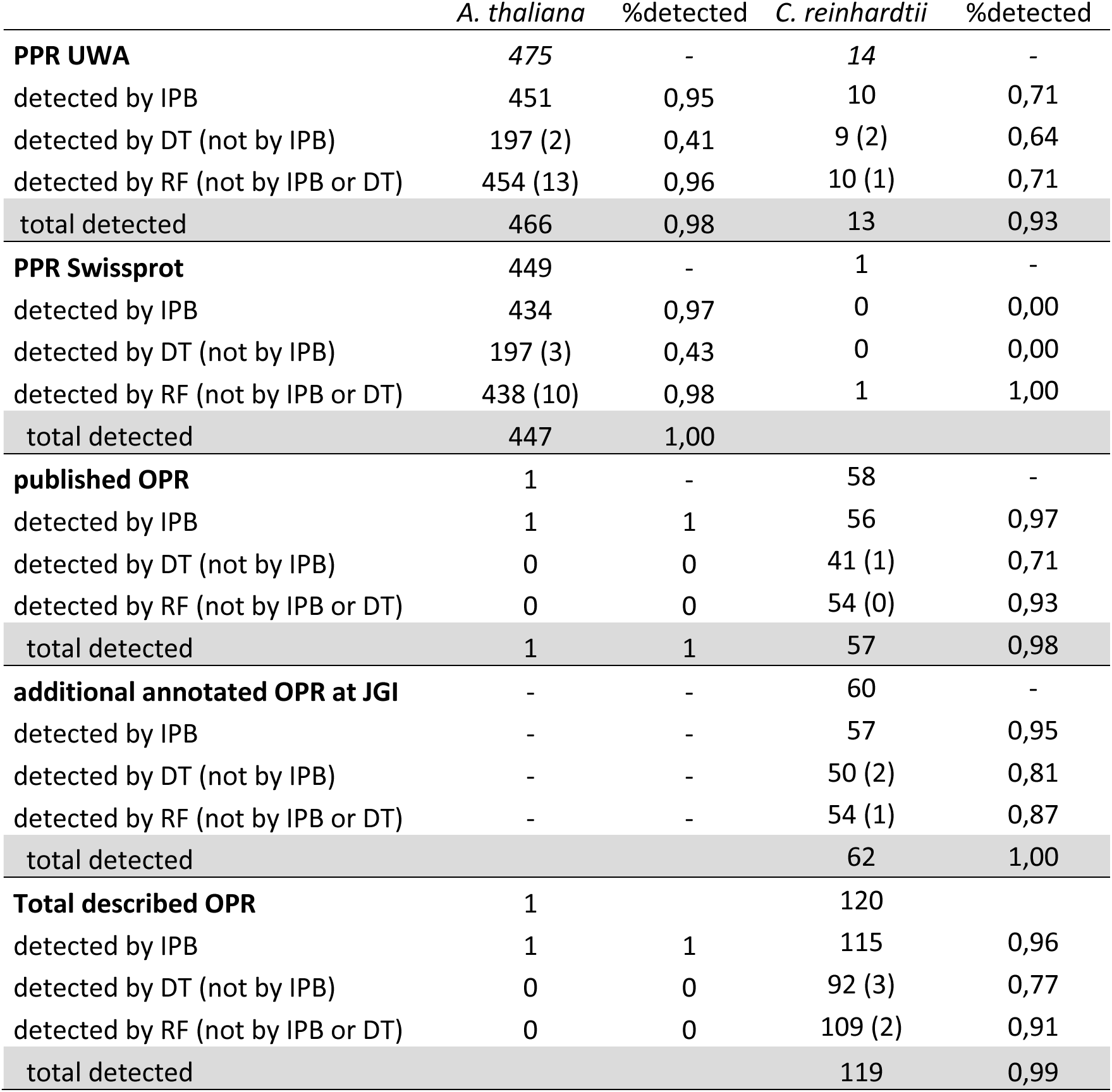
Number of known PPR and OPR detected by the three methods, IPB, DT and RF in A. thaliana and C. reinhardtii. PPR annotated at Swissprot are all the proteins with the term pentatricopeptide or PPR in their names. 3 PPR are annotated by Swissprot but absent from UWA: At1g06143 (PPR15), At1g19525 (PPR51) and At2g13420 (PP150). 33 PPR are unique to UWA. Note that among the PPR list at UWA, 24 are annotated as TPR in Trembl. This is expected as the TPR and PPR are evolutionary related.

|  | *A. thaliana* | %detected | *C. reinhardtii* | %detected |
| --- | --- | --- | --- | --- |
| **PPR UWA** | 475 | - | 14 | - |
| detected by IPB | 451 | 0,95 | 10 | 0,71 |
| detected by DT (not by IPB) | 197 (2) | 0,41 | 9 (2) | 0,64 |
| detected by RF (not by IPB or DT) | 454 (13) | 0,96 | 10 (1) | 0,71 |
| total detected | 466 | 0,98 | 13 | 0,93 |
| **PPR Swissprot** | 449 | - | 1 | - |
| detected by IPB | 434 | 0,97 | 0 | 0,00 |
| detected by DT (not by IPB) | 197 (3) | 0,43 | 0 | 0,00 |
| detected by RF (not by IPB or DT) | 438 (10) | 0,98 | 1 | 1,00 |
| total detected | 447 | 1,00 |  |  |
| **published OPR** | 1 | - | 58 | - |
| detected by IPB | 1 | 1 | 56 | 0,97 |
| detected by DT (not by IPB) | 0 | 0 | 41 (1) | 0,71 |
| detected by RF (not by IPB or DT) | 0 | 0 | 54 (0) | 0,93 |
| total detected | 1 | 1 | 57 | 0,98 |
| **additional annotated OPR at JGI** | - | - | 60 | - |
| detected by IPB | - | - | 57 | 0,95 |
| detected by DT (not by IPB) | - | - | 50 (2) | 0,81 |
| detected by RF (not by IPB or DT) | - | - | 54 (1) | 0,87 |
| total detected |  |  | 62 | 1,00 |
| **Total described OPR** | 1 |  | 120 |  |
| detected by IPB | 1 | 1 | 115 | 0,96 |
| detected by DT (not by IPB) | 0 | 0 | 92 (3) | 0,77 |
| detected by RF (not by IPB or DT) | 0 | 0 | 109 (2) | 0,91 |
| total detected |  |  | 119 | 0,99 |

**Table 3.**
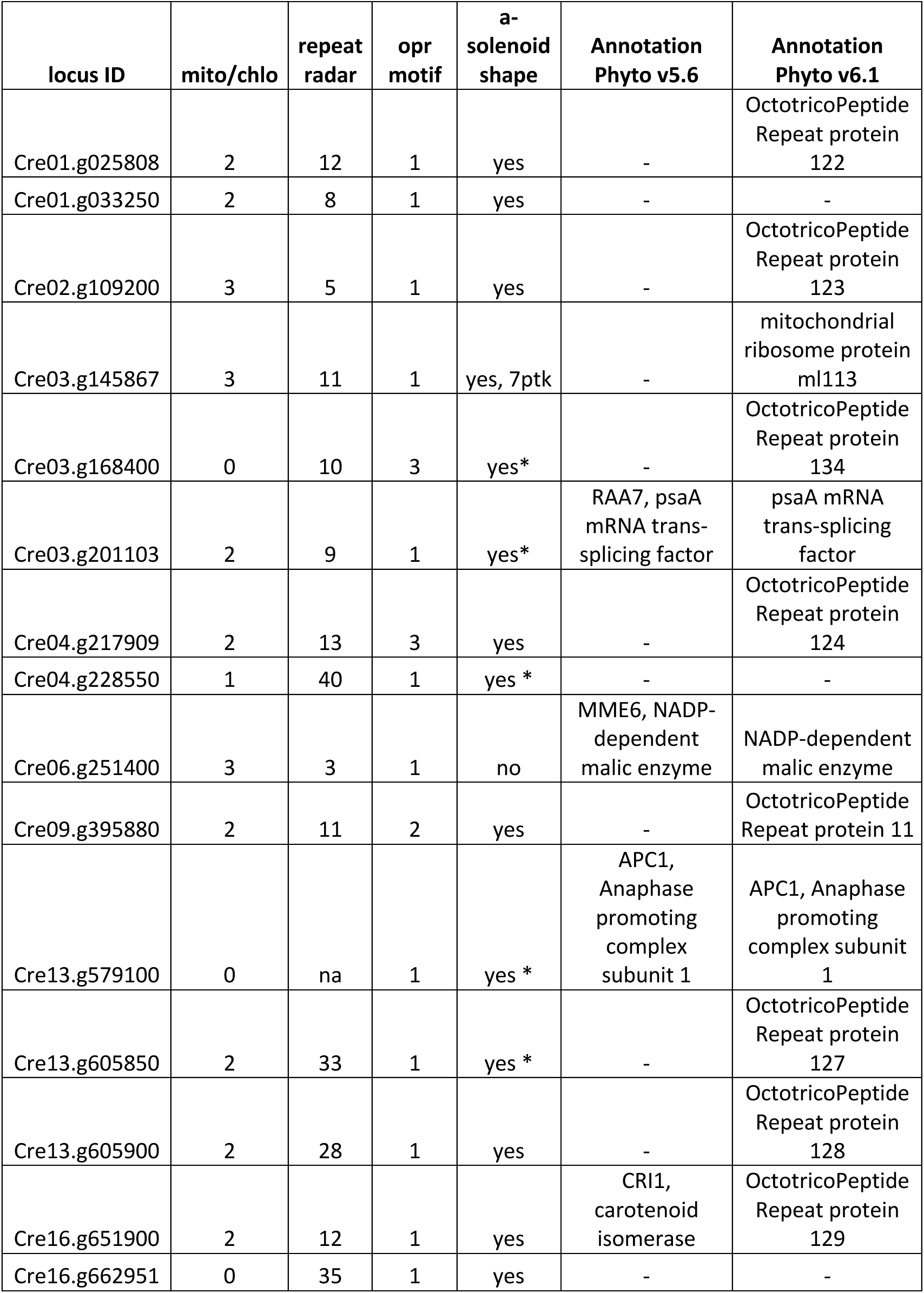
Candidates retrieved by IPB unpublished and unreferenced as OPR with at least 2 motifs (OPR or repeatradar). locus ID: locus identifier in the v5.6b annotation. targ: number of predictions for chloroplast/mitochondrial targeting; radar repeat: number of repeats predicted by RADAR. OPR motif: number of OPR motif found by IPB. a-solenoid shape: Yes/No (AF2 predictions). When an experimental 3D structure is available the PDB accession number is given. * indicates average pLDDT quality scores (< 0.5) for the AF2 predictions.

#### Decision Tree

In the *A. thaliana* proteome, DT retrieved only 197 PPR, 2 of which were not retrieved by IPB (Table 2). DT also found 3 out of the 8 TPR annotated in SwissProt in *A. thaliana*. In the *C. reinhardtii* proteome, 41/50 of the published/additional OPRs were retrieved, all but one (Cre09.g392579) were not retrieved by IPB; DT also retrieved 9 of the 14 PPR present in the *C. reinhardtii* proteome.

#### Random forest

In the *A. thaliana* proteome, the RF procedure retrieved 454 PPR, 13 of which were not retrieved, by either IPB or by DT (Table 2). In *C. reinhardtii*, it retrieved 54/54 of the published/additional OPR. Raa3 and Ncl18, the sole published OPRs that were not retrieved by IPB, were retrieved in 3 and 571 over 1000 RF iterations, respectively. RF also retrieved an additional OPR annotated at JGI (Cre07.g347500, OPR31) that was not retrieved by IPB nor by DT (Table 2).

As shown in Table 2, altogether a same subset of 197 PPR candidates in *A. thaliana* and 91 OPR candidates in *C. reinhardtii* are identified by the three procedures.

By cumulating IPB, DT and RF results, 98% and 99% of the PPR and OPR are detected, respectively.

### New OPR candidates

No new PPR have been identified neither in *A. thaliana* nor in *C. reinhardtii* with IPB, but 15 new OPR have been identified, which were not annotated as such in the v5.6 version of genome annotation in *C. reinhardtii* (Table 3). Among them, 11 are *pto* candidates. Only three of these new OPR candidates have a gene name in the v5.6 annotation available at Phytozome13: i) Cri1 (Cre16.g651900), with 11 repeatsRADAR, annotated as a carotenoid isomerase, adopts an α- solenoid shape predicted by AlphaFold2, (hereafter named AF2pred.); ii) Mme6 (Cre06.g251400) with 3 repeatsRADAR is likely a false positive since its AF2pred 3D structure of low-quality displays α-helices not adopting an α-solenoid shape. (iii) Raa7 (Cre03.g201103), the psaA mRNA trans-splicing factor (Lefebvre-Legendre et al. 2016), with 7 repeatsRADAR. Although other trans-splicing factors Raa1, Raa8 and Rat2 contain OPR motifs (Kück and Schmitt 2021), Raa7 has not been described as an OPR. Its AF2pred. 3D structure although of average plDDT quality score below 0.5, displays pairs of α-helices and could likely fold into an α-solenoid conformation. The 8 other *pto* candidates are uncharacterized and adopt an α-solenoid shape according to AF2pred. Only 2 of the *pto* candidates have more than one OPR motif, but they all contain at least 3 repeats predicted by RADAR (repeatsRADAR). Thus, some OPR motifs have diverged upon recognition by profile similarity, compared to the motifs selected to initiate the IPB procedure (Figure S2), but they are still detected by RADAR.

Note that Cre03.g145867 is part of the large subunit of the mitoribosome (PDB entry 7pkt, chain M).

### New OTAF candidates

Because of the low recall of DT and the low specificity of RF on OPR and PPR *from C. reinhardtii* and *A. thaliana*, we considered the *pto* candidates identified by both DT and RF (*pto* DTRF) as the most robust newly identified OTAF candidates. All candidates retrieved by IPB with 2 OPR/PPR motifs and all *pto* DTRF candidates are listed in Table S4.

Among the 45 *pto* DTRF candidates found in *A. thaliana*, 5 are PPR UWA that were not detected by IPB. Among the 189 *pto* DTRF candidates in *C. reinhardtii*, there is one OPR (OPR14) and one PPR (PPR11) that were not detected by IPB (Table S4). To estimate the quality of the DTRF candidates, they were classified according to their 3D structure predictions (Table S5) in 12 selected proteomes: 3 representative Embryophyta species among Streptophyta (the Bryophyta *Physcomitrium patens,* the Tracheophyta *A. thaliana* and the Marchantiophyta *Marchantia polymorpha*), 3 representatives Chlorophyta species (*Chlorella sorokiniana* in the Trebouxiophyceae, *Ostreococcus tauri* in the Mamiellophyceae and *C. reinhardtii* in Chlorophyceae) and the proteomes of the sole Glaucophyta species and the reference proteomes of the 5 Rhodophyta species. 66% of the *pto* DTRF candidates in those 12 representative proteomes have an α-solenoid structure predicted by AF2 with good quality. If candidates with regions of low-quality predicted structure (plDDT < 0.5) containing only α-helices and disordered regions are considered to potentially adopt an α−solenoïd shape, this number raises to 77% (Table S5).

### Functional annotation of OTAF candidates

Functional annotation of the *pto* DTRF candidates was performed with InterProScan against the PFAM database (v37.4) (Table S6). More than half of the *pto* DTRF candidates have a PFAM domain (1719/3300, 51.1%). Table 4 lists the annotations found in more than 1% of the candidates with a detected PFAM domain. Because InterPro (IPR) annotations summarize results of different databases, IPR annotations are given instead of PFAM domain description to avoid redundancy. The most frequently found IPR annotations for *pto* DTRF candidates are the MYND-type zinc finger domains (169), Ankyrin repeats (63) and Armadillo repeats (53) Note that there are also 129 *pto* DTRF candidates containing PPR/TPR related repeats motifs that were not detected by IPB with the PPR motifs (Table 4). The vast majority (99%) of the PPR candidates have described PPR domains and associated DYW and E domains (Table 4). Because the OPR motif is not described in protein domain databases, the vast majority of OPR candidates lacks characterization. Only 219 out of the 1067 OPR candidates have a PFAM domain (Table 4), 163 of which having a RAP domain, including 10 NCL OPR-RAP tandemly duplicated in *C. reinhardtii* (Boulouis et al. 2015). Note that according to Uniprot annotations, many OPR candidates are composed of disordered regions and regions with compositional biases, either towards acid and basic or polar residues.

**Table 4.**
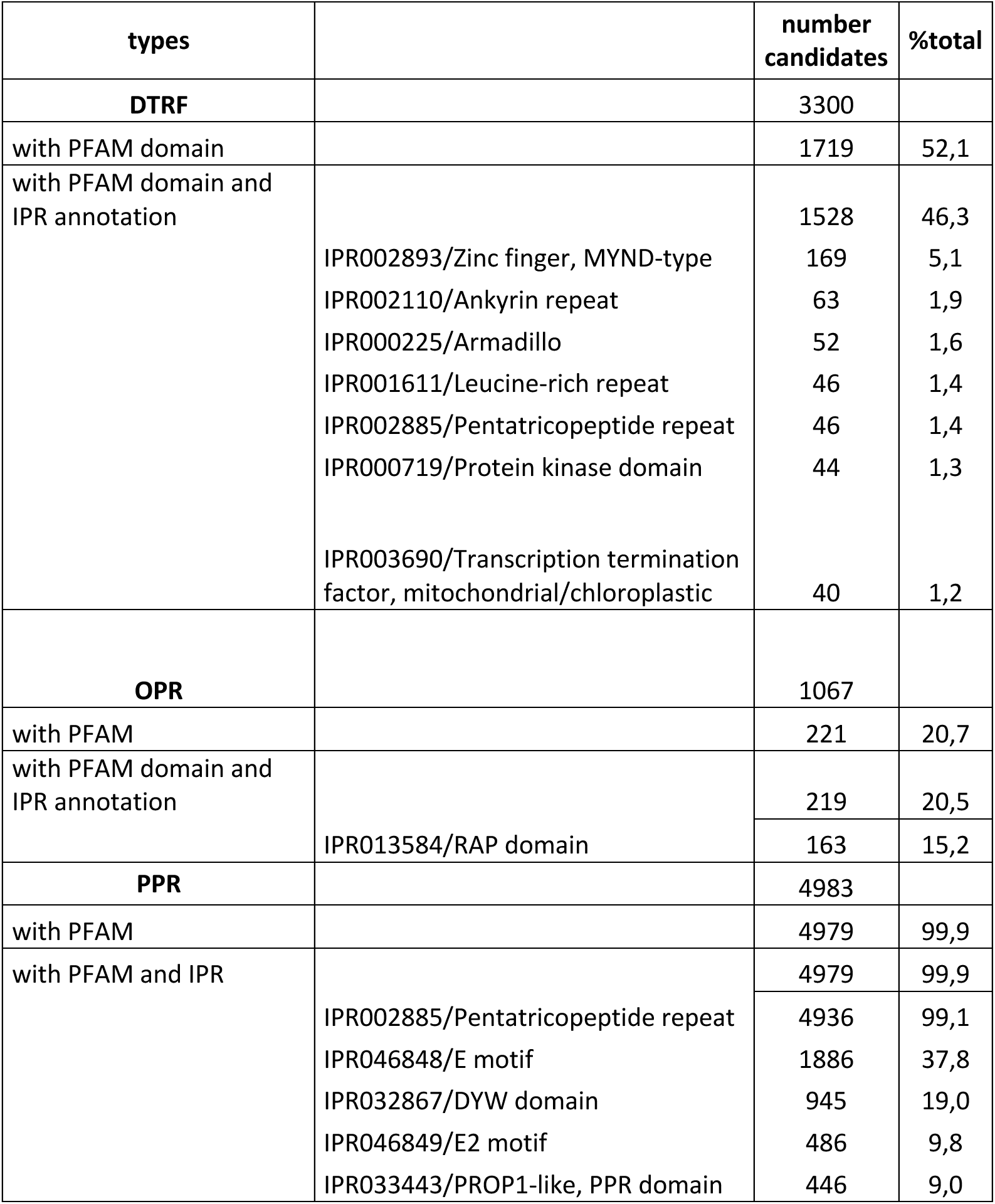
The number of candidates with a detected PFAM motif are given, as well as the subgroup of those candidates with an IPR annotation. To avoid redundancy, the most frequently found annotations are the IPR ones instead of PFAM descriptions.

| types |  | number<br>candidates | %total |
| --- | --- | --- | --- |
| **DTRF** |  | 3300 |  |
| with PFAM domain |  | 1719 | 52,1 |
| with PFAM domain and<br>IPR annotation |  | 1528 | 46,3 |
|  | IPR002893/Zinc finger, MYND-type | 169 | 5,1 |
|  | IPR002110/Ankyrin repeat | 63 | 1,9 |
|  | IPR000225/Armadillo | 52 | 1,6 |
|  | IPR001611/Leucine-rich repeat | 46 | 1,4 |
|  | IPR002885/Pentatricopeptide repeat | 46 | 1,4 |
|  | IPR000719/Protein kinase domain | 44 | 1,3 |
|  | IPR003690/Transcription termination<br>factor, mitochondrial/chloroplastic | 40 | 1,2 |
| **OPR** |  | 1067 |  |
| with PFAM |  | 221 | 20,7 |
| with PFAM domain and<br>IPR annotation |  | 219 | 20,5 |
|  | IPR013584/RAP domain | 163 | 15,2 |
| **PPR** |  | 4983 |  |
| with PFAM |  | 4979 | 99,9 |
| with PFAM and IPR |  | 4979 | 99,9 |
|  | IPR002885/Pentatricopeptide repeat | 4936 | 99,1 |
|  | IPR046848/E motif | 1886 | 37,8 |
|  | IPR032867/DYW domain | 945 | 19,0 |
|  | IPR046849/E2 motif | 486 | 9,8 |
|  | IPR033443/PROP1-like, PPR domain | 446 | 9,0 |

### Distribution of OPR, PPR and new OTAF candidates across Archaeplastida

Figure 2 shows the distribution of the 6050 candidates retrieved by IPB with at least 2 OPR/PPR motifs (IPB2r candidates) and 3300 *pto* DTRF candidates in the 43 studied proteomes. PPR are present in all phyla. There are less than 10 PPR per species in Rhodophyta, Glaucophyta and Chlorophyta and several hundred copies per Streptophyta species, except in the two early-diverging Streptophytes *Mesostigma viride* and *Chara braunii*.

**Figure 2:**
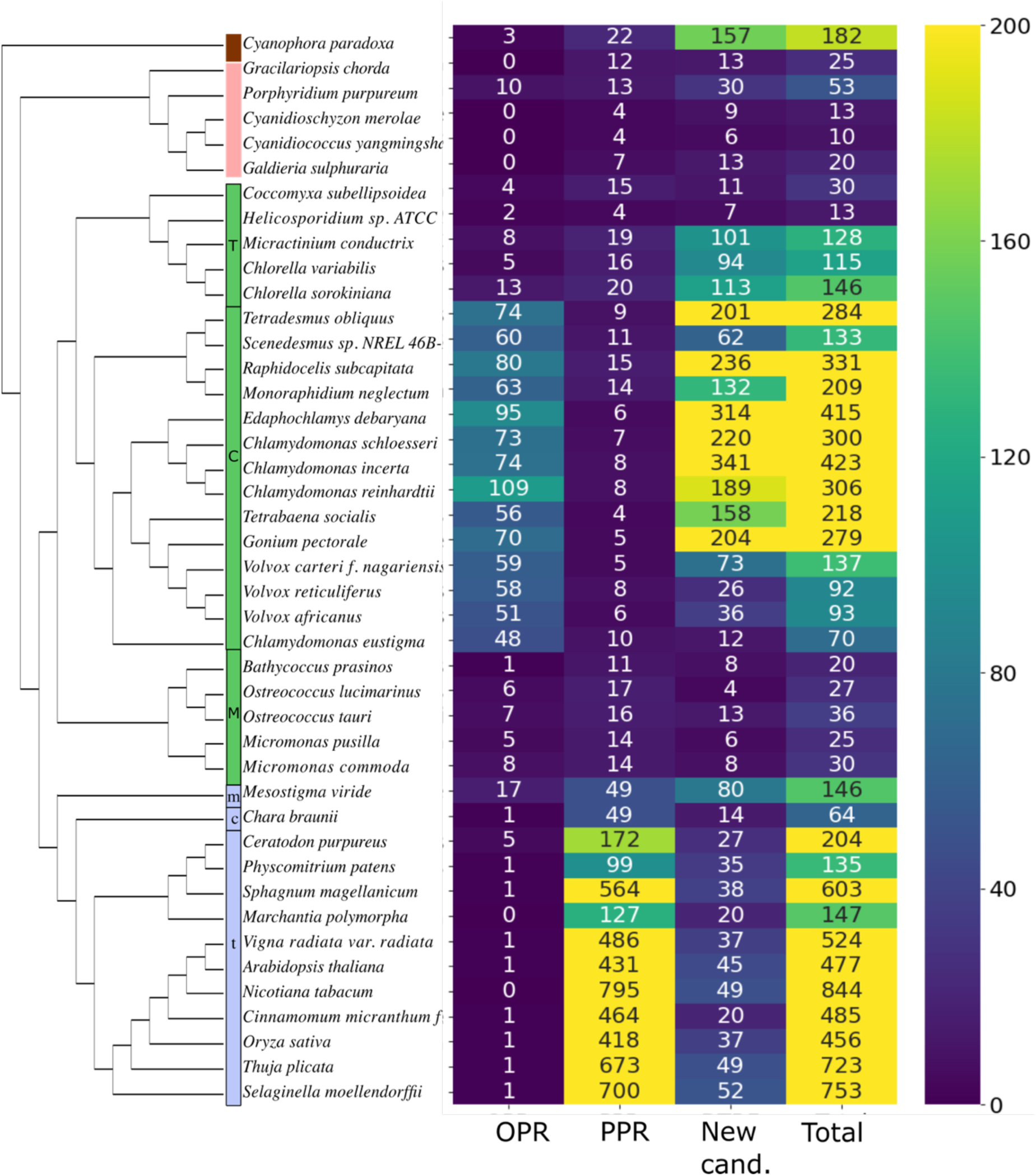
Candidates detected by IPB (with at least 2 OPR or PPR motifs, whatever the number of predictions to be addressed to organelles) and new *pto* OTAF candidates detected by DT and RF across the 43 proteomes of Archaeplastida species, classified according to their NCBI taxonomy and reference phylogenies (see Methods). The last column indicates the total number of retrieved candidates (IPB+DTRF). The color column indicates the different phylum: Glaucophyta (brown), Rhodophyta (red), Chlorophyta (green) and Streptophyta (blue). Sub-clades are indicated within Chlorophyata (T: Trebouxiophyceae, C: Chlorophyceae, M: Mamiellophyceae) and Streptophyta (m: Mesostigmatales, c: Charophyceae, t: Tracheophyta). The whole list of candidates is available in Table S4.

OPR are the most abundant in Chlorophyta, with at least 10 copies, and no less than 48 OPR copies per species in Chlorophyceae, but very few are found in other phyla, except in the early diverging unicellular green alga *Mesostigma viride* (Streptophyta), which contains 17 OPR candidates and in *P. purpureum* which contains 10 OPR candidates. In *P. purpureum*, OPR are automatically annotated as Tbc2 translation factor, chloroplastic, except A0A5J4Z7V5 that remains uncharacterized. If the encoded proteins have the same function than in *C. reinhardtii*, this would suggest that the PsbC mRNA encoding the CP43 subunit of the photosystem II is particularly regulated (Auchincloss et al. 2002). The distribution of *pto* DTRF candidates, as for OPR and PPR, is highly contrasted in the different lineages. For example, there are 157 *pto* DTRF candidates in *Cyanophora paradoxa* compared to only dozens of PPR and 3 OPR. On the contrary, the situation in Rhodophyta is more balanced with only twice as many *pto* DTRF candidates as PPR. Among the 13 DTRF candidates in *Gracilariopsis chorda*, two are annotated as Mbb1, the PsbB mRNA maturation factor. This is also the case of 3 DTRF candidates in *P. purpureum*. While in Streptophyta there are 38 new candidates per species in average, the most contrasted distributions of new candidates are in Chlorophyceae (ranging from 4 in *Ostreococcus lucimarinus* to 341 in *Chlamydomonas incerta*) (Figure 2).

### Potential composite OPR-RAP/PPR candidates

In *Chlorella sorokiniana and* in *Chlorella variabilis,* there are 2 *pto* orthologous candidates A0A2P6U0N6 _CHLSO and E1ZLH2_CHLVA (see below, *Origin of candidates OTAF*) in which IPB detected 4 OPR motifs as well as 1 PPR motif (Figure 3). The sequence logos of the detected OPR and PPR motifs are given in Figure 3B), in which the conserved residues of the canonical motifs are retrieved (see Figure 7 for canonical motifs). The 3D model of A0A2P6U0N6 _CHLSO in Figure 3C, shows two α-solenoid domains that are predicted with high confidence (AlphaFold2 predicted aligned error close to 0 in Figure 3D). The OPR repeats are part of the first one, followed by a RAP domain (in blue and red respectively in Figure 3E,F). The detected PPR repeat is part of the second α-solenoid region (green in Figure 3E,F). The two regions are separated by the first one by a disordered region of approximately 400 amino acids (yellow region Figure 3F). There is a third ortholog in *Micractinium conductrix* for which the PPR repeat was not detected and 2 additional orthologs in *Chlorella ohadii* and *Chlorella vulgaris*, not considered in the present study, but part of the similarity cluster Uniref50_E1ZLH2 which group sequences sharing at least 50% identity (Figure 3A). The domain organization is conserved in these 6 proteins, as exemplified by the dotplot of A0A2P6U0N6 _CHLSO and E1ZLH2_CHLVA (Figure 3G). They would be the first described composite OPR-RAP/PPR candidates. Note that there are much more α-helices in those candidates than the detected OPR motifs, but of which have been detected by RADAR (around 20 repeats in each). This again shows that some PPR/OPR motifs could have diverged upon recognition by profile similarity, compared to the motifs selected to initiate the IPB procedure (Figure S2).

**Figure 3.**
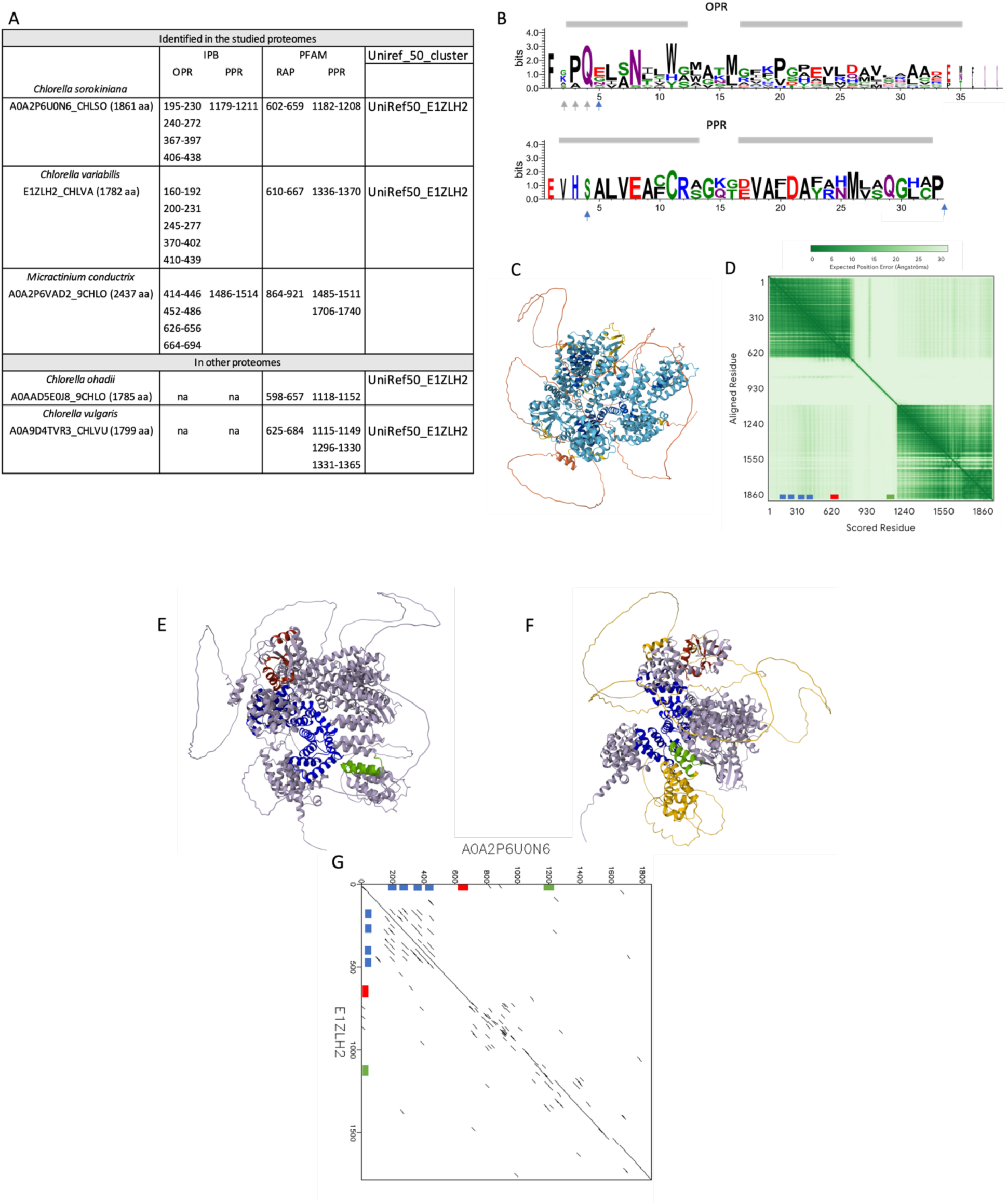
Structure of the composite OPR-RAP/PPR. **A)** Coordinates of the OPR and PPR motifs detected by IPB, and of the PFAM domains detected by InterProScan in 3 candidates identified in this study and the 2 orthologs found in other proteomes according to the Uniref50 clustering. **B)** Sequence logos of the 13 OPR motifs and 2 PPR motifs detected by IPB in the 3 detected orthologs. The positions expected to be determinant for RNA recognition are indicated in blue as in (Jarrige, 2019; Barkan and Small, 2014). Note that the 35^th^ key position is absent from the PPR logo. For OPR, positions indicated in grey are also expected to contribute to RNA recognition. The two putative helices of the AF2pred. of the OPR and PPR sequence logos are shaded in grey. **C)** AF2pred. of A0A2P6U0N6_CHLSO colored according to pLDDT quality score presented in different orientations. **D)** Expected position error of the AF2pred. in C. **E)** and **F)** Different orientations of the structure in C. Repeats detected by IPB are in blue (OPR) and green (PPR) and the RAP domain in red. The region between the RAP domain and the PPR repeat is colored in yellow in panel F. **G)** Dotplot of A0A2P6U0N6 against against E1ZLH2 (made with Dotmatcher at https://www.ebi.ac.uk/jdispatcher). In D and G, approximate positions of the OPR and PPR motifs, and the RAP domain are indicated by blue, green and red respectively

### Origin of candidates OTAF

To date the dynamics of expansion and reduction of the PPR, OPR and DTRF candidates, they were clustered into protein families by MCL clustering (I=50) on their pairwise similarity network (Figure S3). In the present study, we choose to focus on families composed of proteins that did not undergo domain rearrangements since their common ancestor and hence probably conserved similar functions (sequence similarity hits over more than 70% of the length of both aligned proteins). For clarity, the clusters of candidates in the sole 12 representative proteomes are given in Figure 4 and the complete clustering is provided in Table S7. Each family was classified according to the distribution of its members into species-specific, phyla-specific or inter-phyla family (Figure 3). There are 68% (6373/9350) of candidates clustered with at least another candidate, and the majority of them (5113) are distributed into 274 PPR families, 58 DTRF families and 37 OPR families of more than 3 members. There are 25/9 DTRF candidates clustered within PPR/OPR protein families, thus corresponding to remote PPR/OPR homologs, as mentioned earlier. In agreement with PPR frequency and the similarity network (Figure S3), the three largest protein families comprise a total of 3080 PPR candidates in all 4 phyla studied. In contrast, the largest DTRF and OPR families are restricted to Chlorophytes, and are much smaller (78 and 32 homologs for DTRF and OPR respectively).

**Figure 4.**
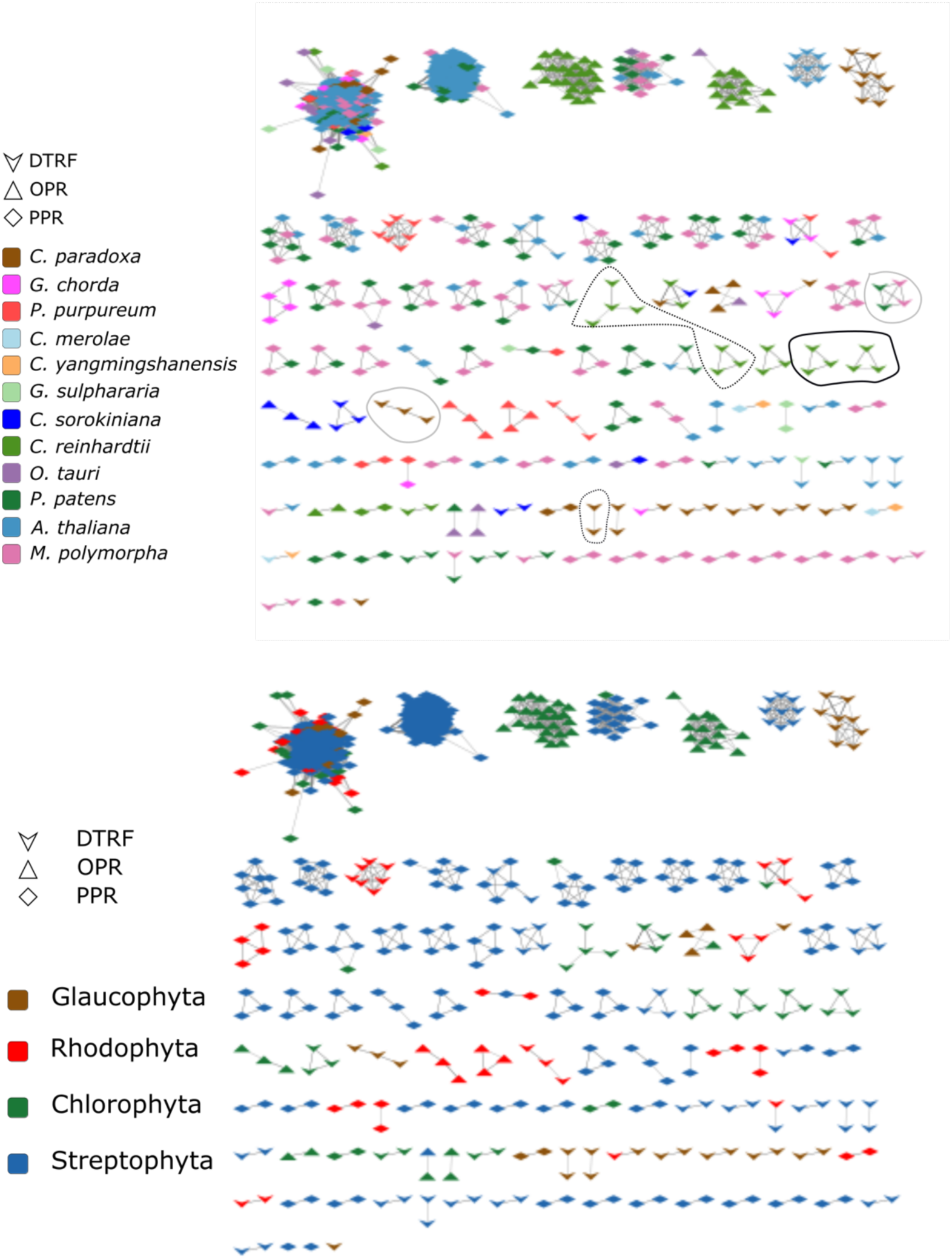
**A.** Clusters of IPB and *pto* DTRF candidates found in Glaucophytes and Rhodophytes, as well as in 3 representatives in both Streptophytes and Chlorophytes. *Pto* candidates are colored per species and their shape indicates the detection method (legend box). Pto candidates linked by grey edges are in the same cluster and share significant similarity (E-value < 10e-6) over at least 70% of their length. Clusters of DTRF candidates containing different type of repeats are surrounded: TPR repeats (PFAM domains, grey), Ankyrin repeats (dotted black, no PFAM domain but CATH superfamily) and Mynd-type clusters (black, no PFAM domain but CATH superfamily). **B.** Same as in A, with candidates colored according to their taxonomical group.

Compared to OPR and DTRF candidates, PPR are the most frequently clustered into inter-phyla families (57%) and into Streptophyta-specific families (33%). PPR would thus be the most ancient families, or at least the most conserved ones, with the lowest frequency of singletons (4,8%). At the opposite, there is only one OPR family with members in more than one phylum. The majority (89%) are either within Chlorophyta-specific families (52%) or as singletons in Chlorophyta (41%), suggesting that OPR emerged more recently or evolved faster than PPR.

There are 35 families of DTRF candidates with at least 5 members across the 43 studied proteomes (TableS7). 70% of DTRF candidates are singletons, suggesting that most of the families among DTRF candidates emerged more recently or evolved faster than both OPR and PPR (Table S7). The largest families are restricted to Chlorophyta, but 3 large ones are shared between the phyla: one of 11 ankyrin repeats shared by 10 species in all phyla except in Rhodophyta, one of 9 peptidases shared by 8 species in all phyla but Glaucophyta and on of 8 TPR undetected by IPB in Chlorophyta and Rhodophyta. Higher inflation (I) values for MCL clustering would split the largest PPR families into several smaller ones, thus inverting the proportion of inter-phyla families (from 55% at I=50 to 24% at I=100) and phyla-specific families (from 33% at I=50 to 59% at I=100, Table S7), but it has only very minor effect on OPR and DTRF clustering. In consequence, the general tendencies of the comparative analysis of PPR, OPR and DTRF families are robust: PPR are the most shared between phyla and have the fewest proportion of singletons.

Species-specific families, likely reflecting recent duplications or fast evolving candidates, are observed in all species, with the highest duplication rates in the moss *Selaginella moellendorffii* (Streptophyta) for PPR, in *Edaphochlamys debaryana* (Chlorophyta) for OPR and in *Cyanophora paradoxa* for *pto* DTRF candidates (Table S7). There are 11 species-specific expansions of at least 5 DTRF paralogs found in each phylum. In those families, we note the high frequency of proteins containing Zinc fingers MYND-type (Table 5). Note that Zinc fingers MYND-type domains have been found only in Chlorophyta (Table S6).

**Table 5.**
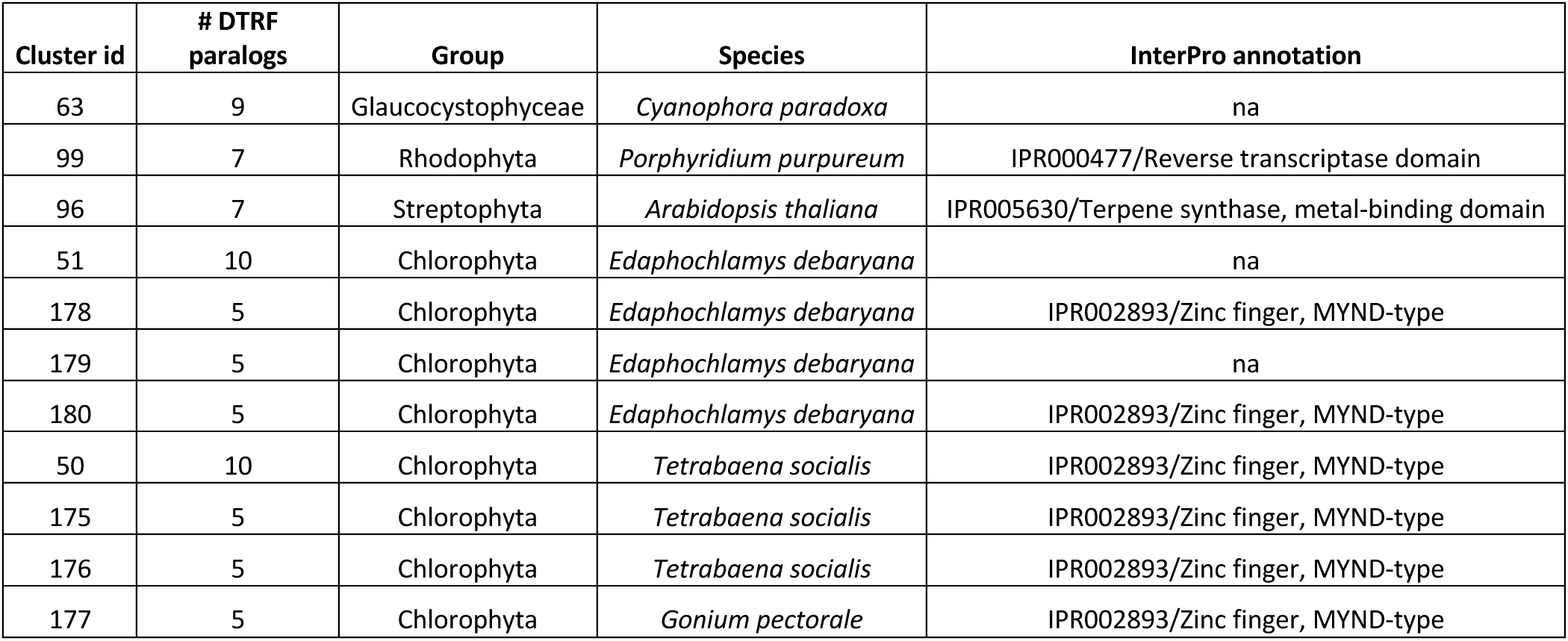
Species-specific protein families

There is a notable species-specific expansion of interest in *C. reinhardtii*. It contains 7 paralogs tandemly duplicated on chromosome 10, comprised of 2 pto DTRF candidates, one other DTRF candidate and 4 non-*pto* RF candidates (Figure 5A). The 7 paralogs in that duplicated region have an average pairwise identity of 68% over at least 85% of their sequence and adopt an α-solenoid fold (Figure 5B). No PFAM domain is detected within these candidates. There are also 3 homologs of that family in *C. incerta* and 2 homologs in *C. schloesseri* (cluster 94, Table S7.1). In each of these two genomes, the paralogs are not tandemly duplicated, in agreement with an expansion restricted to *C. reinhardtii*.

**Figure 5.**
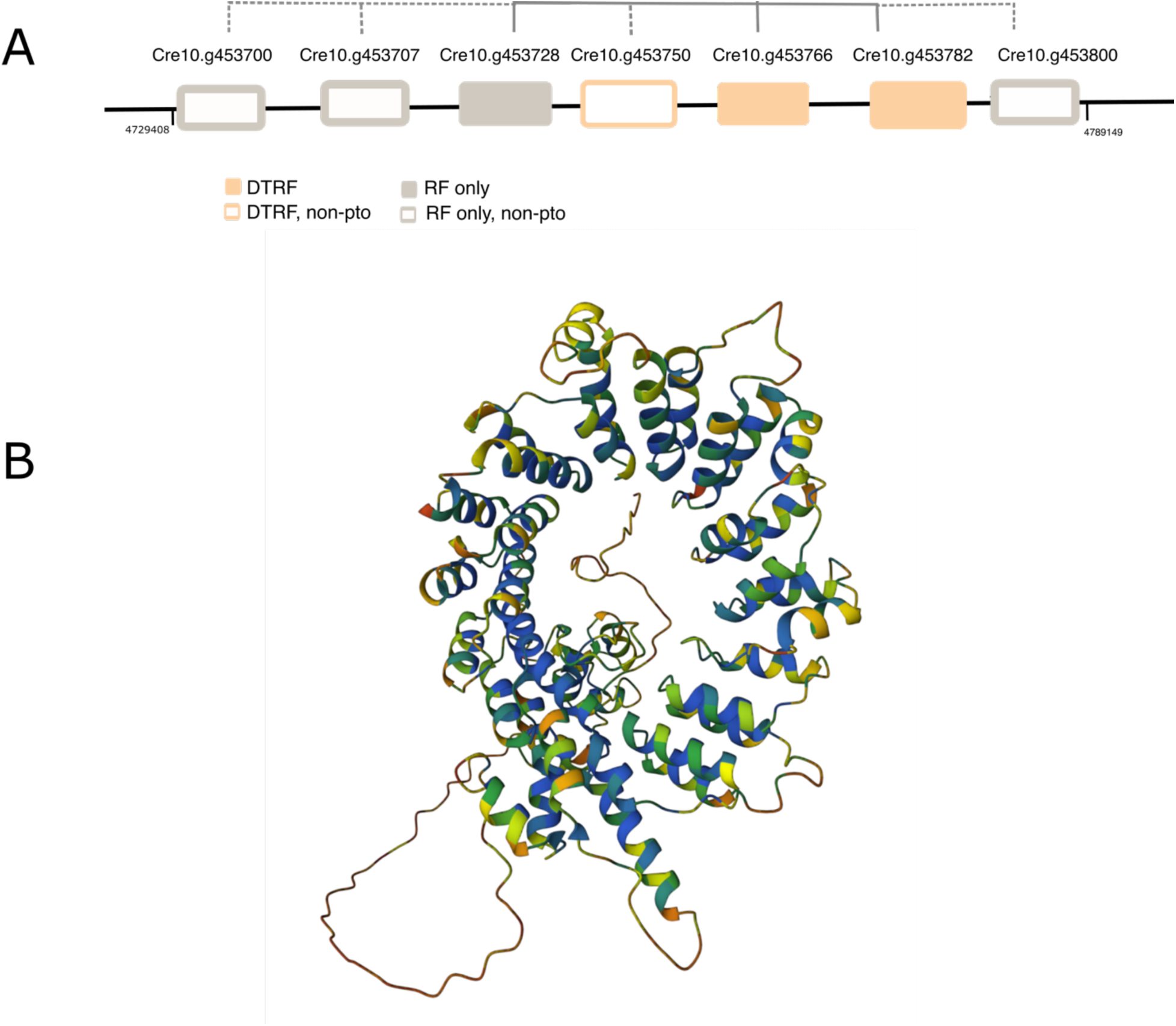
**A.** Tandems of DTRF candidates on chromosome 10 of *C. reinhardtii*. Solid grey lines relate proteins in the same cluster of *pto* candidates (Fig. 4). Dashed grey lines relate non-*pto* to *pto* candidates. **b)** AF2pred of Cre10.g453800. All proteins are classified in the CATH Superfamily 1.25.40.20 termed ankyrin repeat-containing domain (InterPro IPR036770) but they do not contain any PFAM Ankyrin or ankyrin-related repeats.

### Molecular and physico-chemical properties of OTAFs candidates

#### Number of repeats

The number of OPR and PPR motifs detected by IPB is given in Figure 6A. Half of the PPR candidates found by IPB have more than 7 motifs and in *C. reinhardtii*, the number of OPR motifs per candidates follows a bimodal distribution with a pic at 2 and the other at 7 motifs per proteins. As already mentioned, some OPR and PPR motifs diverged upon recognition but are detected by RADAR (Figure S2). This explains why the PPR motif per protein detected by IPB is lower than the median number (9-10) of PPR motifs described among P-type PPR proteins found in land plants (Cheng et al. 2016), and of course lower than the median number of PPR motifs (12) among all PPR UWA.

**Figure 6.**
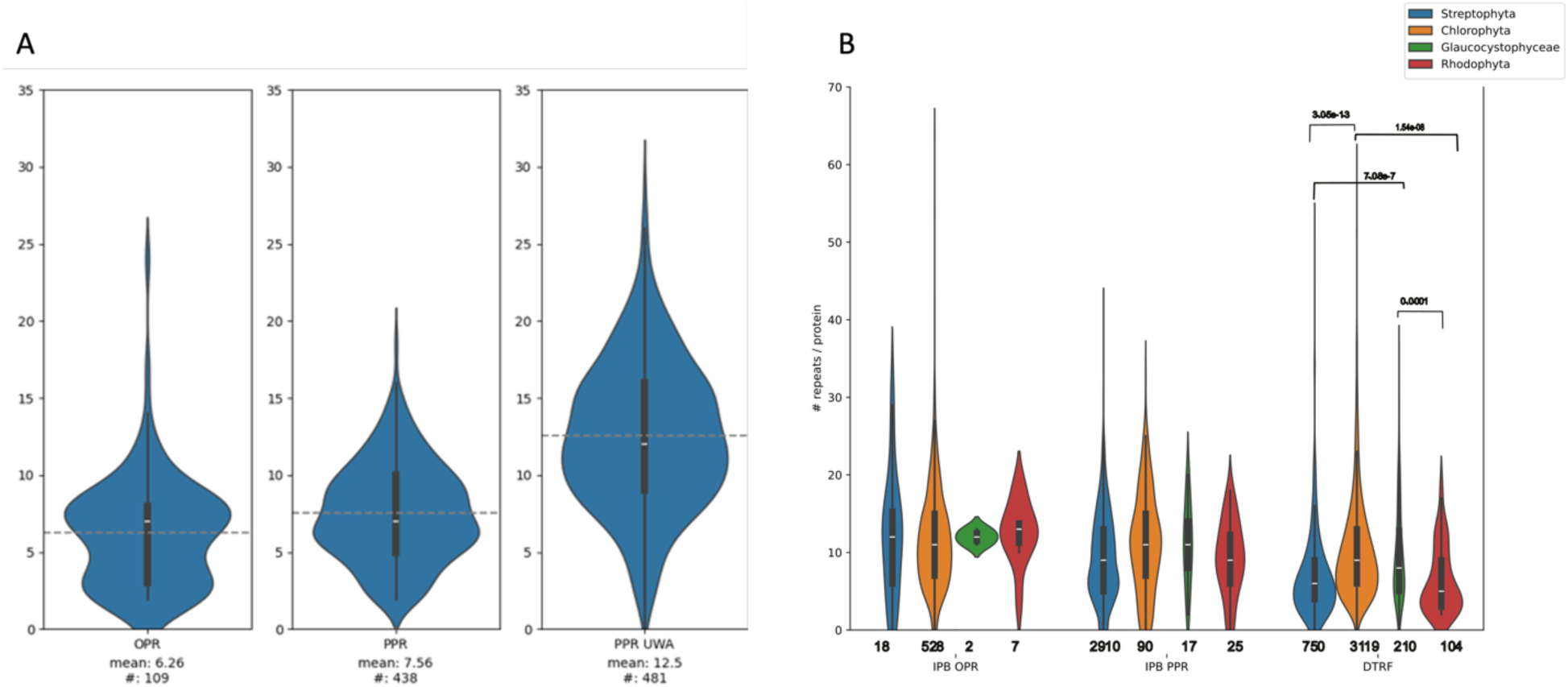
**A.** Left and middle panels: Distribution of the number of OPR/PPR motifs in IPB candidates with at least 2 detected motifs in *A. thaliana* and *C. reinhardtii*. Right panel: Distribution of the number of PPR motifs per PPR UWA. **B** Distribution of repeats per protein detected by RADAR for the IPB and pto DTRF candidates in the 4 taxons studied. The number of *pto* candidates is indicated below and significant p-values (post-hoc Tukey test, p-value < 0.05) above the violinplots.

The number of repeats detected either by IBP or RADAR in each type of OTAF candidates among the 4 phyla is shown in Figure 6B. The number of OPR and PPR repeats per *pto* candidates is similar in each phylum. Inversely, the number of repeatsRADAR in the *pto* DTRF candidates vary between phylae, with the highest number being in Chlorophyta. This suggests, in line with the duplication/loss rates described above (Table S7) a higher diversity of protein families and a higher propension to form repeats in Chlorophyta.

#### Clustering of OPR and PPR motifs

The similarity networks between OPR and PPR motifs found in the 12 representative proteomes is shown in Figure S3A. While most of the PPR motifs form a unique network, OPR form one big network with distinct sub-networks by species and 6 smaller networks of *C. reinhardtii* OPR motifs. Motifs were clustered (see Methods) and classified as species-specific, phyla-specific or shared among different phyla (Table S9). Motifs within a clustermotifs share on average 55% identity while motifs between two different clustersmotifs share on average 0.1% identity (Figure S4B).

Most of the cluster motifs are phyla-specific. In agreement with the distribution of the protein families, PPR clustersmotifs are mostly found in Streptophyta proteins and OPR clustersmotifs in Chlorophyta proteins. Species-specific clusters and singletons also exist (36% for PPR motifs and 12% for OPR motifs). This suggests a heterogeneity of evolutionary rates among clusters. PPR motifs are globally more conserved, but there are also rapidly evolving PPR motifs within Streptophyta species and rapidly evolving OPR motifs in Chlorophyta. Fewer inter-phyla clustermotifs are found for PPR and OPR (*ca.* 4% and 3%, respectively). The existence of PPR/OPR clusters motifs with no *C. reinhardtii*/*A. thaliana* members confirms the ability of IPB to retrieve motifs diverged from the ones used for the first iteration step.

On average, all the motifs within a given clustermotifs belong to different proteins. In other words, the different motifs that compose an OPR/PPR protein are part of different clustersmotifs. Since our study spans 43 proteomes, this could reveal greater similarity between homologous motifs in homologous OPR/PPR than between the motifs that make up a given protein. Alternatively, this could reveal a positional bias of the different clustersmotifs along the protein. To investigate this hypothesis, we determined the order of appearance of each clustermotifs along the proteins (repeat relative position). For this analysis, a MCL clustering was performed on the similarity network of the 706 OPR motifs and the 3346 PPR motifs found in *C. reinhardtii* and *A. thaliana* only, to exclude the orthologs of the 41 other species . For each cluster, only one representative among the different paralogs of a given protein family were considered. The left panel of Figure 7A shows the heatmaps of the frequency distribution of clustersmotifs (> 4 motifs) along PPR proteins. For example, the PPR motifs of clustermotif 1 are found as the first repeat of PPR proteins less than 10% of the time, and at similar frequencies at other relative positions along the protein. Thus, the frequency distribution of clustermotif1 is not biased. Only three PPR clustermotifs present a strong positional bias (at least 80% of the motifs at a given relative position) along PPR candidates. PPR cluster 12 and 18 are mostly found as the first repeat and cluster 29 as the second repeat along PPR proteins. The logos of different type of PPR motifs are shown on the right panel of Figure 7A. The logo of all PPR motifs described at UWA (UWA logo) is given on the top. It is mostly similar to the logo of the P motif, which is the most frequent PPR motif. The strong similarity of the logo for all motif found by IPB to the UWA logo confirms the performance of IPB. Interestingly, the conserved Glycine at the 15^th^ position of the UWA logo is substituted in the clusters 12 and 29 with positional bias : (G->S) within clustermotifs 12, biased at the first position and (G->K) for clustermotifs 29. Clustermotifs 29 also have a substitution (G->Y) at the 31^st^ position of the UWA logo. Position 15^th^ lies within the linker between the two α-helices and position 31^st^ lies just after the second α-helix. These substitutions might thus provide different flexible properties to the repeat. Clustersmotif with positional biases have also more hydrophobic residues in the first part of the second helix that might also provide different structural conformations, like the higher global hydrophobicity of the first positions within clustermotifs 12. Also note that the variability of the residues at the first key positions for mRNA binding specificity is lost (N only) for cluster 29, suggesting that these biased motifs repeat binds to a Cytosine according to the code proposed in (Barkan and Small 2014) if the second position is the 38^th^ one in cluster 29, but if it is the 35^th^ one, the nucleic acid specificity is not provided by the code. Importantly, note that the biased clusters do not match the sequence logo of the variant of the PPR motif L and S (Barkan and Small 2014). Interestingly, the linker region of the consensus PPR motif from the 17^th^ residue up to the 30^th^ residue in the second helix contains mainly hydrophobic amino acids, with 5 negatively charged ones evenly distributed. The negative charge of this region is completely lost in clustermotif 12, in which the corresponding positions contains either positively charged residues (2) or neutral ones (3). In clustermotif 18, the negativity also decreases, as positively charged residues at position 13^th^ and 16^th^ are the most frequently observed.

Figure 7B shows the heatmap and logos for OPR motifs. Clustermotif 10 are mostly found as the first repeat along OPR proteins and cluster 11 are mostly found as the second repeat. The linker region in the consensus OPR motif (16^th^ to 20^th^ position) is overall positively charged (+1), as well as in the cluster with no positional bias (+2), while in the cluster with positional bias towards the first or the second position within the proteins, the overall charge becomes negative (-1). Also, in the clustermotif 11 biased towards the second position, the overall hydrophobicity is lost.

#### Physico-chemical properties

All OTAF candidates were described with the variables used in the RF procedure. Figure 8 shows the principal component analysis (PCA) of the 1551 candidates within the set of selected 12 proteomes used to evaluate the quality of AF2pred. of DTRF candidates. Overall, candidates of a given phyla are close together in the two-dimensional space defined by the first principal component (PC1) and PC2 (Figure 8A). They differ mainly along the PC1 axis, with OPR proteins on the right (Figure 8C) and PPR proteins on the left (Figure 8D). OPR and PPR thus differ mainly by their hydrophobic and steric properties (Figure S5), which is also visible on their sequence logos (Figure 7). Some DTRF candidates co-localize in the same area as OPR/PPR candidates retrieved by IPB, indicating that the similarity-free detection procedures succeed at identifying the defined properties of known OTAFs (Figure 8B). Other DTRF candidates explore a wider space through the upper-right part of the graph, away from OPR and PPR, indicating higher amphipathic properties (Figure S4), that could also reflect contrasting structural properties and thus functions.

**Figure 7.**
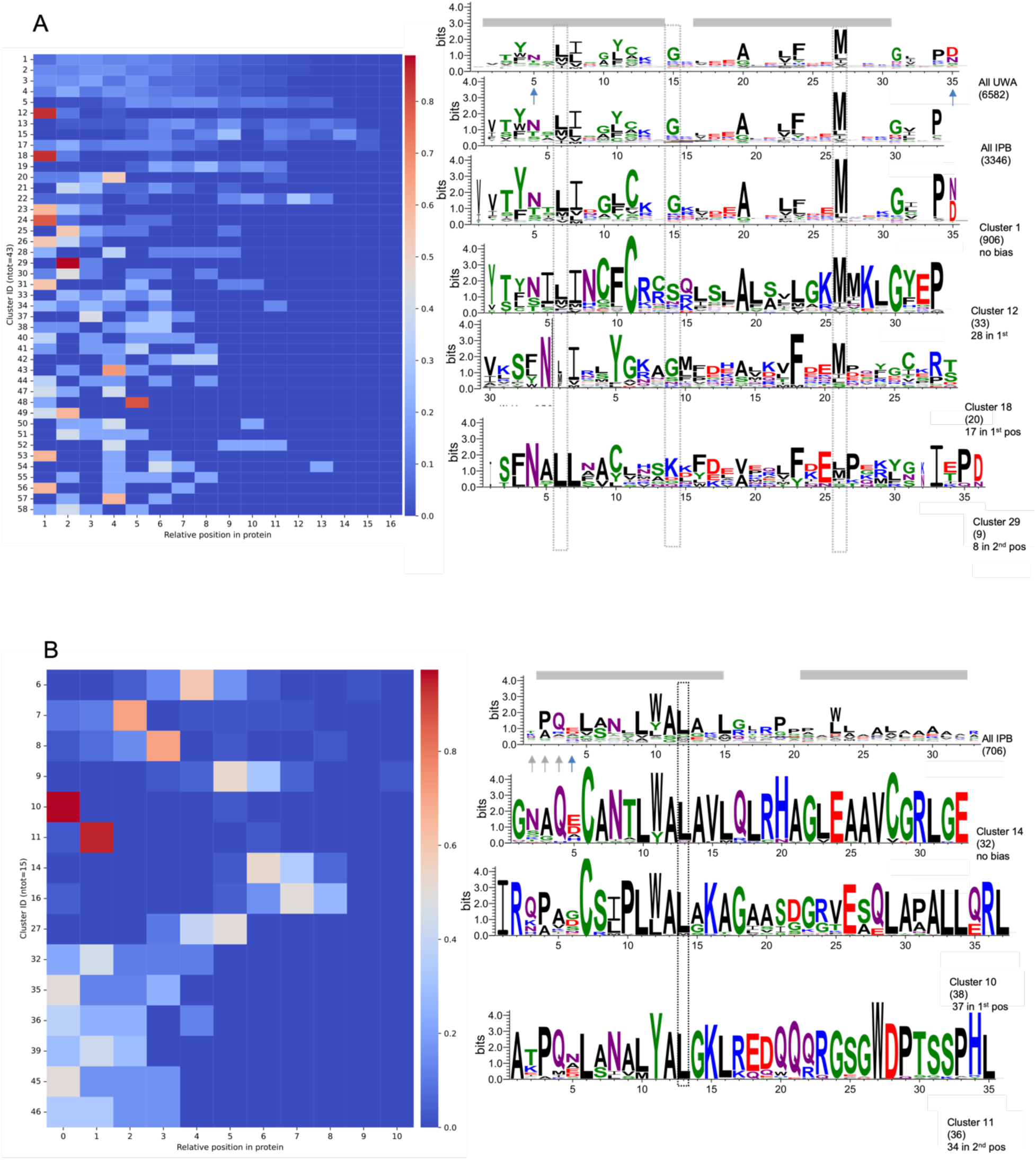
Distribution of clustermotifs. A. PPR motifs. Left panel, heatmap of the frequencies of each clustermotifs (motifs > 4) according to their relative position along the PPR protein. Right panel, logos of remarkable clustermotifs. On top, logos of PPR motifs described at UWA and all PPR motifs found by IPB. The amino acid positions are those obtained from the multiple alignment and for PPR motifs at UWA it coincides to those proposed based on structural data in (Yin et al. 2013). All logos are aligned on the Methionine at the 27^th^ position of PPR motifs at UWA. Conserved residues important for the structure of the motifs are boxed on Figure 7A. The N-terminus of first helix of clustermotif 18 (30th to 35th position) was found after the second helix in the logo. In the figure, those residues have been moved at the beginning of the motif. As in (Barkan and Small 2014), the two helices of the PPR motifs are shaded in grey and the positions shown to be the primary determinants of RNA-binding specificity are indicated by arrows. A selection of conserved positions in the three regions of the PPR motif are boxed in dotted dark. For each logo, the number of motifs in the cluster is indicated in parenthesis. When the distribution is biased, the occurrence of motif at the biased position is given. Amino acids are colored according to their chemistry (green, polar; purple, neutral; blue, basic; black, hydrophobic). There are on average 9, 9, 5 and 8 PPR motifs per proteins in clusters 1, 12, 29 and 18 respectively. B. Same for OPR motifs. All logos are aligned on the 12^th^ position of the logo of all detected OPR motifs, boxed in dotted dark. The two putative helices of the AF2p OPR sequence logo are shaded in grey and the position expected to be the primary determinant of RNA-binding specificity as in (Jarrige 2019) is indicated by a blue arrow. The three positions immediately upstream, indicated by grey arrows, are also likely contributing to the RNA recognition. There is are on average 8 OPR motifs per proteins in clusters 6, 10 and 11.

**Figure 8.**
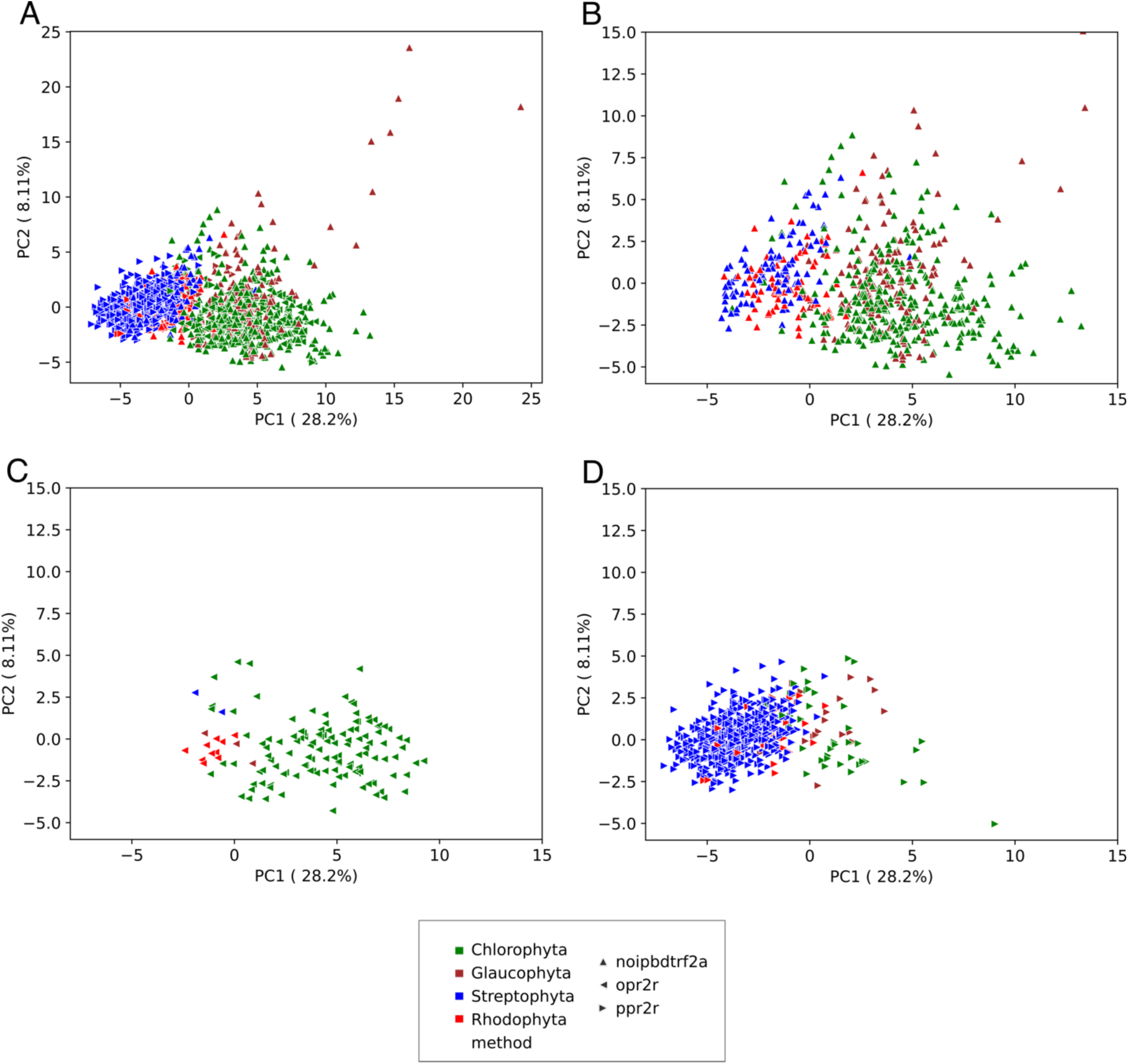
Principal component analysis of candidates with at least 2 OPR/PPR motifs and DTRF candidates in 12 representative proteomes of Archaeplastida described by properties used by the RF procedure. **A** All candidates along principal components 1 and 2. **B** DTRF candidates (same PCA as in A). **C**. OPR candidates (same PCA as in A). **D**. PPR candidates (same PCA as in A). Correlation circles are given in Figure S5.

## Discussion

### Critical assessment of IPB

IPB, the iterative similarity-based procedure efficiently retrieves the OPR and PPR previously identified in *C. reinhardtii* and *A. thaliana*. It even identified new remote homologs in these two species, seven of which have been annotated as new OPR (Cattelin 2023) in version 6.1 of *C. reinhardtii* (Craig et al. 2023). Starting from motifs found in a single species, the iterative process allowed us to capture specific signatures in diverse taxonomic groups to identify OPR and PPR homologs across Archaeplastida species. On average we found less PPR motifs by PPR candidates than those reported at UWA. This discrepancy is mainly due to the fact that we used a very limited number of PPR P-motif in IPB. In contrast, PPRFinder, used to establish the PPR UWA catalog (Gutmann et al. 2020), searches a dozen of profiles of PPR sub-motifs that were expertly defined combining analyses of both sequence and structural similarity based on the analysis of 41 land plants proteomes (Cheng et al. 2016). Despite its simplicity, IPB proved to be an efficient approach for detection of conserved motif at a broad taxonomy scale, with no *a priori* knowledge on the motif diversity and its tridimensional structure. Its sensibility may be improved by increasing the number of motifs used to build the original profiles; by evaluating the quality of multiple alignments obtained by several methods with metrics as proposed in (Mirarab and Warnow 2011) and also by evaluating the quality of the similarity clusters obtained by alternatives to MCL for clustering (such as the Louvain algorithm).

### Critical assessment of DT and RF

The DT procedure is very specific but poorly sensitive (0.34), which explains why it failed to detect several of the known OTAFs. In contrast to DT, the RF procedure is very sensitive but lacks specificity on the real datasets. It could be improved by using additional tools for single residue 2D predictions such as NetSurfP (Høie et al. 2022) for repeat *ab initio* detection and α−solenoid structures. As for any supervised classifier, RF candidates are strongly determined by the training set. There are also many possibilities to improve the RF procedure, starting by testing other protein descriptors. Some attempts to develop machine learning procedures have been already made for PPR discovery, based on protein features extracted from PPR as a positive training set and non-PPR as negative training sets as in (Qu et al. 2019; Feng et al. 2021; Zhao et al. 2021b) but the performance of these methods have been estimated only on the training sets: they were never used to search PPR in complete proteomes, and therefore it is not known how they compare to profile PPR-based methods in such tasks and how it copes with taxonomical bias. Indeed, deep learning approaches using natural language processing models and neural networks models need also be tested. However, when taken together, DT and RF identified new *pto* α−solenoid candidates (DTRF), of yet unknown OTAFs families. We showed that hydrophobic and steric constraints on the same face of a α-helix (Figure S1) have a crucial role for the RF classification. Such properties could reflect amphipathic constraints and hence the amphipathic nature of the helices, necessary to allow the formation of a hydrophilic cavity within the α- solenoid structure, in which the positively charged mRNA could bind. Based on our estimation that 66% of *pto* DTRF candidates have an actual α-solenoid shape. Note that this evaluation could be automated in the future by using pipelines like the recently developed SOLeNNoID (Nikov et al. 2025) to detect solenoid residues into protein 3D structures.

### Completed catalog of OPR and PPR across Archaeplastida

PPR are the most conserved OTAFs between phyla and have the fewest proportion of singletons, compared to OPR and DTRF candidates. Massive expansions of PPR candidates occurred in Tracheophyta, which is probably linked to the expansion of mRNA editing in this lineage among land plants (Gutmann et al. 2020). The relative paucity of PPR in green microalgae could be associated to the absence of RNA editing in Chlamydomonas (Gallaher et al. 2018), as well as, to our knowledge, in Rhodophyta and Glaucophyta. If such events were to be identified in those clades, they would be mediated by players other than PPRs, in agreement with the proposal that eukaryotic editing was probably acquired multiple times from ancestral bacterial toxin deaminase (Iyer et al. 2011). IBP allowed us to provide the first extensive catalog of OPR protein across Archaeplastida. They are mainly found in Chlorophyta, with few members in Streptophyta, but also in Rhodphyta and Glaucophyta, confirming the trends already found by comparing micro-algae and land plants. The HPR described in (Hillebrand et al. 2018) are recognized by IPB as harboring OPR motifs, confirming that HPR proteins belong to the OPR family (TableS10).

### A potential bias in the distribution of the repeated motifs along the OPR and PPR proteins

The clustering of the motifs provided by IPB also pointed to a potential bias in the distribution of the clusters of PPR and OPR motifs along a protein sequence, with no periodicity of occurrence along the protein, like for the PLS subfamily of PPR proteins, composed of repetitions of P, L and S consecutives motifs (Cheng et al. 2016). The analysis of the sequence logo of the biased OPR and PPR motifs shows that the overall charge of the linker regions in those motifs (and the second helix for PPR) vary compared to the consensus motif. To our knowledge, such a bias has not been reported previously. It calls for further investigations to assess the contribution of such bias for particular position in mRNA target recognition and/or protein structural properties, such as folding and flexibility. As the linker region lies on the external surface of the α-solenoid, this could play a role for binding of other protein partners.

### Have composite OPR/PPR proteins appeared in an ancestral Trebouxiophyceae ?

Thanks to IPB, we identified 5 orthologous proteins composed of both an OPR-RAP and a PPR α- solenoid domain in Trebouxiophyceae. The fact that those proteins were identified in 5 different species makes the probability of 5 independent sequencing or assembly error quite low. Such composite protein has never been observed before. If they arose by a chromosomal rearrangement in an ancestral Trebouxiophyceae, their conservation suggests that they could have been selected during evolution. Further investigations will be performed to determine if they are real proteins and, if so, to determine their function.

### New α-solenoid OTAFs candidates available for functional characterization across Archaeplastida

There are on average *ca.* one hundred of new α-solenoid OTAFs candidates in Chlorophyta and in the Glaucophyta *Cyanophora paradoxa*, and fewer in Streptophyta and Rhodophyta, with 29 candidates in *A. thaliana* and 19 in *Porphyridium purpureum*. We thus provide a -most probably incomplete- tentative list of new OTAF candidates, for further experimental characterization. As expected, among the *pto* DTRF candidates, some display conserved domains typically defining α−solenoid proteins like armadillo and ankyrin repeats domains. Such proteins have been found in land plant chloroplasts, some likely being involved in plastid gene expression, such as the ankyrin repeat protein Akrp in *A. thaliana* that blocks chloroplast differentiation (Zhang et al. 1992) and an armadillo/β-catenine protein in spinach, found in the chloroplast nucleoids which might be involved in the structural maintenance of chromosomes (Melonek et al. 2012). Interestingly, a significant number of *pto* DTRF candidates of Chlorophyta, have Zinc finger MYND-type domains, which predominantly mediate protein-protein interactions and are found in transcriptional regulators as well as DNA repair proteins (Kamaliyan and Clarke 2024). Also, many DTRF candidates contain intrinsically disordered regions, which interestingly have been found to contribute to single stranded RNA binding of Zinc finger domains, although of another type (Kijima and Kitao 2025). Many other *pto* DTRF candidates have no conserved PFAM domain, but are classified into the CATH Superfamily database, based on domains found in proteins with defined structure, that allows to find more remote similarity. This is the case of the cluster of 7 paralogs tandemly duplicated found in *C. reinhardtii* (Figure 7), containing domains from the superfamilies of ankyrin and pseudo-ankyrin repeats. This cluster might represent a reservoir of new candidates and its experimental characterization is ongoing, by looking for a possible photosynthetic phenotype of the corresponding mutants in *C. reinhardtii*.

### The exploration of α-solenoid OTAFs candidates outside Archaeplastida made possible

Little is known as to how the expression of the organellar genomes is regulated outside of Opistokhonta and Viridiplantae (land plants and green algae). Our analysis showed how contrasted is the global distribution of putative α-solenoid OTAF proteins, with far less OTAFs, either known (OPR/PPR) or unknown (*pto* DTRF) in Rhodophyta and Glaucophyta than in Viridiplantae, suggesting that modes of regulation may vary, possibly with a different balance between transcriptional and post-transcriptional steps in the corresponding organelles. In particular, the RF approach should be instrumental for studies of the largely unexplored chloroplast biology from Rhodophyta and Glaucophyta, but also in other types of photosynthetic eukaryotes, such diatoms or dinoflagellates, which result from secondary or even tertiary endosymbiosis, paving the way for elucidating the rules of the regulation of gene expression in organelle genomes in these complex plastids.

## Materials and methods

### Sequence data

The 43 proteomes from Archaeplastida species were retrieved either at NCBI, Uniprot or JGI websites. The details of the proteomes and their versions are given in Table S1. For proteomes downloaded at JGI, the file of the filtered models was chosen Files/Annotation/Filtered Models ("best")/Proteins. Quality and completeness of proteomes were assessed with BUSCO 5.3.2 (Manni et al. 2021) (Table S1), with protein mode on the Eukaryota and Viridiplantae lineages (eukaryota_odb10, viridiplantae_odb10, 2020-09-10).

As starting points for the motif-based procedure (IPB), we used published and described repeat motifs from 12 known OPR proteins from *C. reinhardtii* (green microalga model species) and from 11 known PPR proteins from *Arabidopsis thaliana* (land plant model species), listed in Table 1.

### Annotations, structural predictions

Proteins annotations were retrieved at Uniprot for all proteins. For *C. reinhardtii*, the annotation v5.6 available at Phytozome, version 13 was also used, as well as the TAIR10 annotation for *A. thaliana*. 3D structure predictions were retrieved at the 1Fold Protein Structure Database or estimated locally with AlphaFold2 (Jumper et al. 2021). PFAM domains were searched with InterProScan (v5.75-106.0) (Jones et al. 2014) against PFAM 3.47 in InterPro 106.0. For each candidate, if a PFAM domains is detected, the IPR annotation is provided. Secondary structure assignment from PDB coordinates was made with STRIDE (Heinig and Frishman 2004).

### Sublocalization prediction

Deeploc v2.0 (Almagro Armenteros et al. 2017), LOCALIZER v1.0.4 (Sperschneider et al. 2017), TargetP-2.0 (Emanuelsson et al. 2007) and WoLFPSort v0.2 (Horton et al. 2007) were used to predict the chloroplast or mitochondrial localization.

### Reference phylogeny

The 43 species from Archaeplastida were selected based on proteome status as provided by the Published Plant Genomes (PPG) database (https://www.plabipd.de/) on June 2020. Reference phylogeny was inferred from PPG and the following articles <u>(Nakamura et al. 2013; Wickett et al</u>. <u>2014; Lemieux et al. 2015; Leliaert et al. 2016; Liu et al. 2019; Nakada et al. 2019; Cheng et al.</u> <u>2019; Hanschen and Starkenburg 2020; Zhao et al. 2021a)</u>, with the help of the NCBI Taxonomy tool.

### Motifs-based similarity procedure

The IPB procedure follows the four steps described below, that we ran iteratively either on OPR or on PPR motifs.

#### Step 1. Pairwise comparison of motifs

First, an all-against-all pairwise comparison of all motifs is performed by BLASTP v2.6.0 (Camacho et al. 2009). We kept only the pairs of motifs whose e-values were lower than a threshold *t* varying according to *X* the size of the data set, as follows: if 10*^n^*^−1^ ≤ *X* ≤ 10*^n^*, then *t* = 10^−*n*^.

#### Step2. Motifs clustering

Second, motifs are clustered with the MCL v.14-137 (Enright et al. 2002) based on the −*log*(*E*−*value*) of the BLASTP hits. Clusters are computed with inflation parameter I values, starting from 1.1 and by increasing it by step of 0.1 until. For each I parameter tested, a multiple alignment of the motifs in a given cluster is performed with MAFFT v7.450 (Katoh and Standley 2013). We choose the clustering with the I value providing a maximum number of clusters with at most 18 and 16 gaps in their multiple alignment, for OPR and PPR motifs respectively. These values correspond to half of an OPR or a PPR motif as we wanted to detect motifs that were similar to the starting motifs over at least half of their sequence and avoid “motif slippage”.

#### Step3. HMM profiles search against proteomes of interest

Third, profiles are built with *hmmbuild* from the HMMer suite, v3.1b2 (Mistry et al. 2013). Each HMM profile was then searched against each of the 44 proteomes with *hmmsearch*. The length of an OPR/PPR motif being about 35/38 amino acids, only newly found sequences whose lengths are -6/+2 the length of an OPR motif, *i.e.* 32/40 amino acids long are kept. Two motifs overlapping by more than 80% are considered identical and the motif boundaries are defined by the minimum overlapping region to reduce the tendency of the motifs from slipping in one side or the other. Two motifs overlapping by less than 20% are considered different and kept without changing their boundaries. Pairs of new overlapping motifs between 20% and 80% are both eliminated, being considered as no longer representative of the typical starting motif.

#### Step5. Selection of candidates on targeting properties

The procedure is iterated restarting from step 1 after adding the new motifs to the initial data set only if at least 1 new *pto* candidate, *i.e.* predicted to be addressed by at least two prediction softwares is retrieved. See “prediction of localization” paragraph for details of the prediction.

#### Step6. Selection of the candidate proteins

The final step consists in selecting only *pto* proteins, i.e. predicted to be addressed either to chloroplast or mitochondria by at least two prediction algorithms (see *Sublocalization prediction*).

### Decision tree (DT) procedure

First of all, proteins with a predicted transmembrane helix after its N-terminal part (corresponding to the targeting peptide and that could erroneously be predicted as a transmembrane helix) were removed. TMHMM v2.0c (Krogh et al. 2001) was used to predict transmembrane helix. The end of the N-terminal part was defined to the 78^th^ residue.

The DT procedure is given in Figure 1, right panel. Parameters for selection along the decision tree have been defined based on a set of validated α-solenoid protein and a set of validated non α-solenoid proteins (see the paragraph *Training set composition*). Table S3 recapitulates numbers of retrieved candidates, precision, accuracy and recall for all tested combinations of parameter values. We choose the global combination maximizing precision and recall.

#### Filtering of proteins according to the number of sequence repeats

We selected the proteins in which at least one repeat of at least 29 amino acids is detected by RADAR v1.3 (Heger and Holm 2000). 29 amino acid length gives the best precision score among a series of tested values, from 20 to 40 (Table S3).

#### Detection of α helix pairs

The presence of α-helix pairs was estimated with ard2 (Fournier et al. 2013), which uses a neural network procedure with a training set containing X-ray resolved α-helix pairs and gives a score for each amino acid position to be a linker between two α helices.

In order to keep only proteins with multiple pairs of α-helices, we constructed and tested dozens of sets of criteria based on the distance between 2 α-helix pairs and the number of linkers between the helices forming a pair (Table S3). We kept only the proteins with at least 4 linkers, with the distance between two consecutive linkers comprised between 32 and 400 amino acids. A given amino acid is considered as a linker if its ard2 score is above 0.15, based on the results obtained on the training set.

#### Protein filtering according to 2D structure predictions

We kept only those proteins for which 65% of the amino acids were predicted in α-helix by S2D v2 (Sormanni et al. 2015), between the two most extreme linkers predicted by ard2. We tested for 50% to 100% and we chose the percentage filter having the best precision score (65%, Table S3). Finally, only *pto* candidates are selected using the same targeting prediction tool as in the IPB procedure.

### Random forest classifier

As for DT, proteins with a predicted transmembrane helix after its N-terminal part are removed. The RF classifier uses the properties also used in the DT procedure: the median of the length of the detected repeats, the number of linkers predicted by ard2 (see above), the median of the distance between two linkers and the probability for an amino acid to be a linker. The proportion of residues in the detected repeats are considered rather than the number of repeats to have a continuous distribution of values. In addition, the RF model uses the proportion of residues with disorder propensity predicted by IUPred3 (Erdős et al. 2021) with score > 0.5 and default parameters, the 20 amino acid frequencies and 36 Auto-Cross Correlation (ACC) values of neighboring amino acids over a window of 4 residues computed for each protein based on the Z-scales amino acid descriptors (Hellberg et al. 1987) as described in (Garrido et al. 2020). The training set was divided in a validation set (10%) never used for learning or testing and the remaining 90%. We then selected the best model using the best_estimator function from gridsearchcv. We evaluated the performances of the best model using best_model.predict on the validation set and it gave an accuracy of 98,61%. The best hyperparameters were 500 estimators, a maximum tree depth of 10, a minimum number of 5 samples required to be at a leaf and all other parameters at default values. The confusion matrix comparing the expected classification results with those obtained from the model prediction (see figure S6) shows that only one protein out of 72 was misclassified.

The RF classification was performed with the bestmodel.predict function from the scikit-learn Python package (version 0.21.2). We used the mean decrease in impurity procedure to determine the importance of each feature in the model. Finally, only *pto* candidates are selected as in IPB and DT.

### Training set composition

The training set for DT and RF procedures comprises 426 known α-solenoids (positive set) and 286 known non α−solenoids (negative set) manually selected based on their 3D structure, resolved experimentally or predicted by AF2. The distributions of the properties used by DT and RF for the training set are given in Figure S7. Note that in the negative controls, proteins that adopt solenoid shape made of β-sheets and α−helices were included in an attempt to increase specificity towards “all-α” solenoids.

### Motifs analysis

Multiple alignments of motifs were obtained by MAFFT v7.450. Positions with more than 80% of gap residues were filtered and used to create the logo with the standalone version of WebLogo 3.7.9 (Crooks et al. 2004).

### Clustering

Clustering was performed with MCL v.14-137 (Enright et al. 2002) based on the −*log*(*E*−*value*) of the BLASTP hits. For the determination of the Inflation parameter in IPB, see IPB section above. For family reconstruction, the Inflation parameter I values of 20 to 80 were performed. 50 was selected as it is the one were DTRF and OPR families were stable. For the final clustering of OPR and PPR motifs, the Inflation parameter was set to 2 and the cluster was performed on the identity percentage rather than the E-value, on hits sharing similarity over at least 80% of their length. Network visualization was performed with Cytoscape (v3.9.1) (Shannon et al. 2003).

### Statistical analysis

Statistical tests and PCA were performed with the scikit-learn Python package version 0.21.2.

## Acknowledgments

We thank Hédi Soula for fruitful discussions and advice for setting up the machine learning procedure; Richard Lavery for initial discussions on molecular dynamics; Yves Choquet for fruitful discussions on OPR; Olivier Vallon for providing access to the unreleased OPRdb database and for stimulating discussions with CC. This work was supported by funding from the Centre National de la Recherche Scientifique and Sorbonne University to UMR7141; the “LabEx Dynamo” (ANR-LABX-011), the DECRYPTOR grant (CNRS 80prime 2019) and the DecryProtARN grant (CNRS Infiniti 2018).

## Notes

### Competing Interest Statement

The authors have declared no competing interest.

