## Supplementary material for "Searching α−solenoid proteins involved in organellar gene expression": Cattelin_FigSupp.pdf

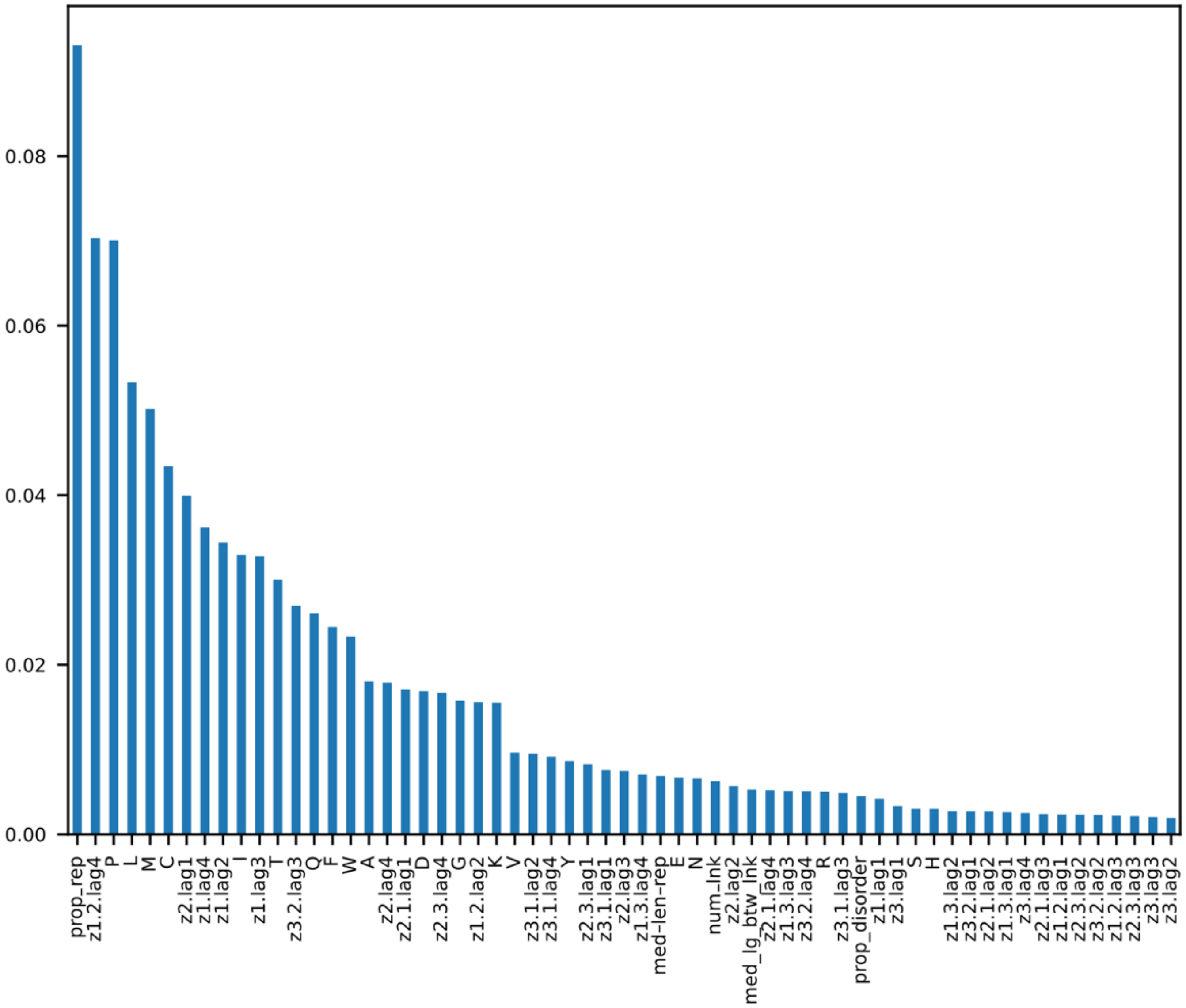


**Figure S1**. Importance of the variables in the RF classifier. Y axis corresponds to the mean decrease in accuracy (MDA). Prop_rep : proportion of amino acids in repetitions, med-len-rep : median length of repetions, num-lnk : number of linkers detected between 2 alpha-helices, med_lg_btw_lnk: median length between linkers. Prop-disorder : proportions of amino acids in intrinsic disorder.

The second most important property is the AAC term z1.2.lag4, thus combining Z1 and Z2. The Z1 scale reflects hydrophobicity and the Z2 scale reflects the steric properties (bulkiness) between residues n and n+4 (lag4). At the level of a $\alpha$-helix of 3.6 amino acids per turn, residue n+4 can be considered as lying on the same face of the a-helix than residue n. Hydrophobic and steric constraints on the same face of a $\alpha$-helix could reflect amphipathic constraints. The sulfur-containing Methionine and Cysteine are considered to help structure stabilization and are also important for redox-regulation.

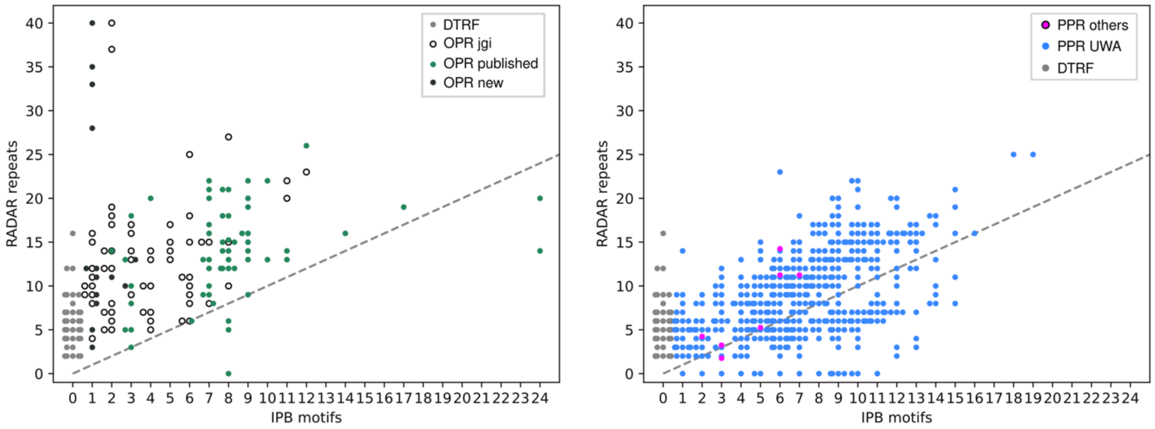


**Figure S2.** Number of OPR/PPR motifs detected by IPB and RADAR repeats in IPB and DTRF candidates found in *C. reinhardtii* (left) and *A. thaliana* (right). For IPB candidates, only those found with OPR/PPR motifs are shown for *C. reinhardtii/ A. thaliana.* The dashed grey line is the diagonal. OPR jgi : annotated OPR at JGI, in addition to the published ones (see Table 2). The jitter option was used to reduce overplotting of the multiple points for a given position.

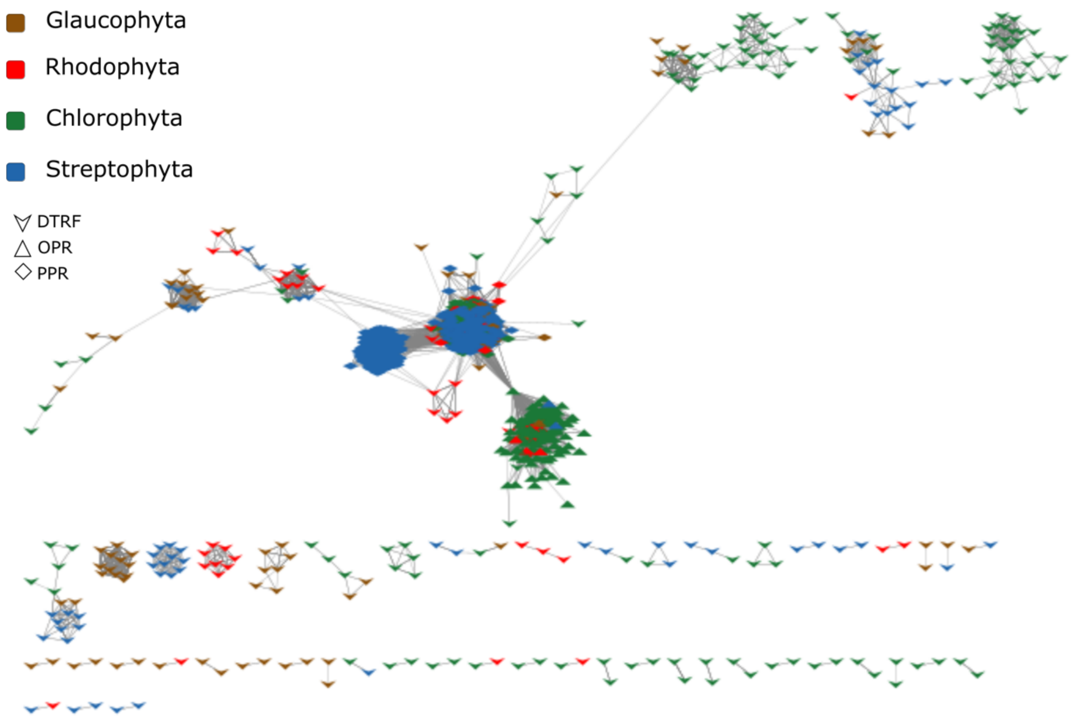


**Figure S3**. Similarity network of the 1551 IPB and *pto* DTRF candidates (only 1212 share Blast hits with E-value < $10^{e-6}$) from the representative proteomes used in Figure 3A. The edge connecting the the big PPR sub-network to the big OPR sub-network is between composite OPR-RAP/PPR candidates (Figure 3).

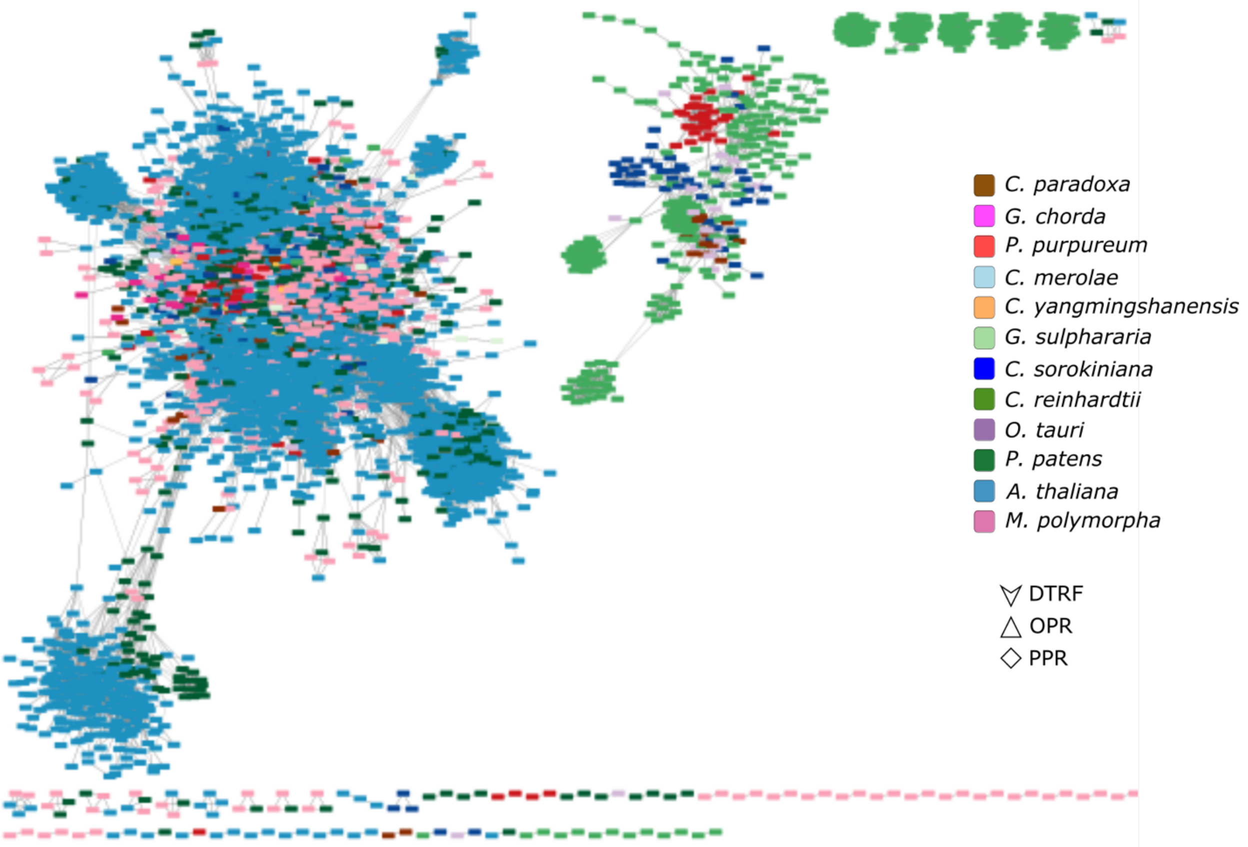


**A**

**B**

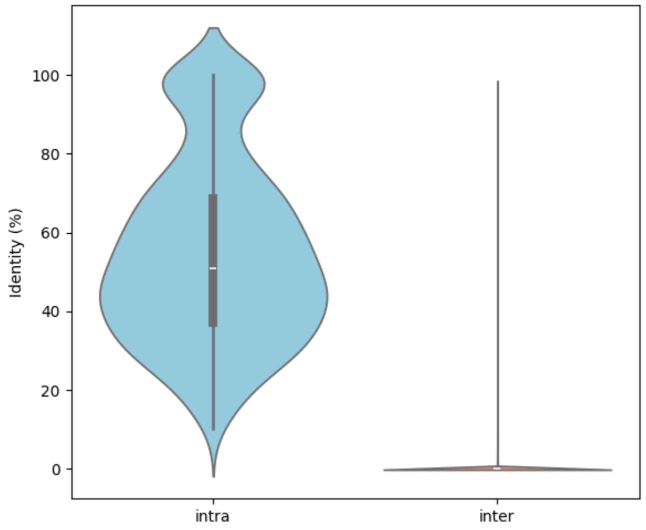


**Figure S4. A**. Similarity network of the 6058 motifs (813 OPR and 5245 PPR) found in the twelve representative proteomes (only 5489 share Blast hits with E-value < 10e-6). Same color legend as Figure 3A. **B**. Identity percentage between motifs within MCL clusters motifs (intra, left) and between motifs from different clusters (inter, right).

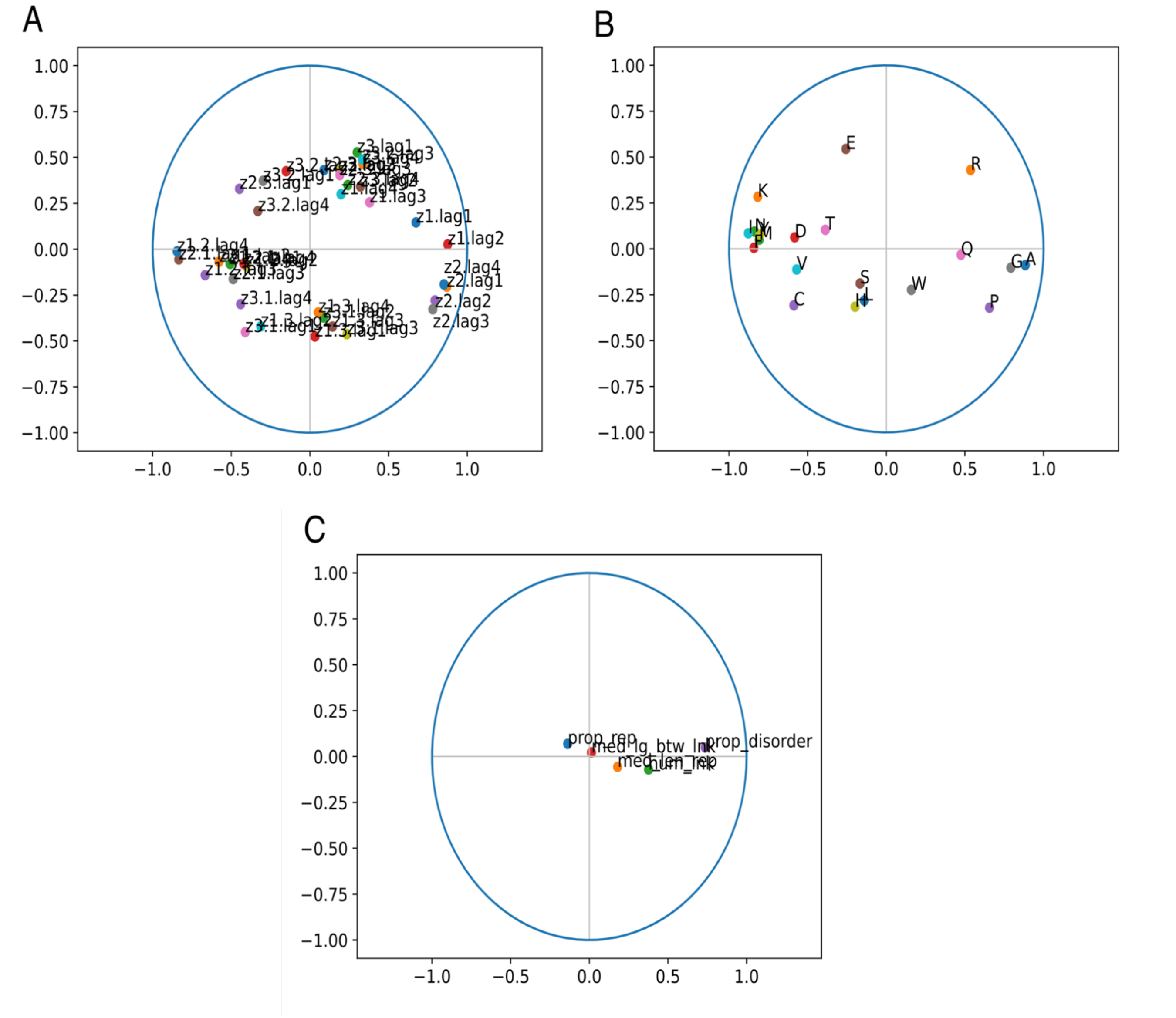


**Figure S5**. Correlation circles of the PCA of Figure 8 for ACC terms (A), amino-acid frequencies (B) and other properties (C). The correlation plots of the PCA are given in Figure S3. The proportion of residues in repeats and the number of linkers between two alpha-helices have only a moderate contribution in the PCA (Figure S5), which can be explained by the fact that these properties were used to select the candidates.

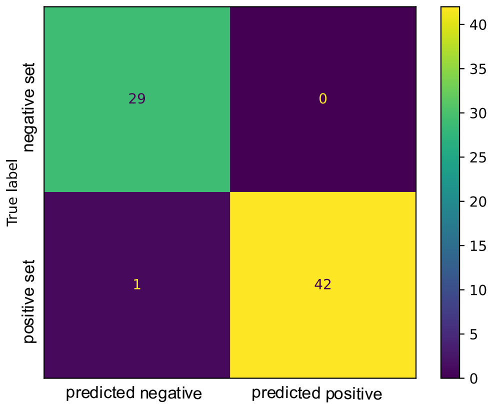


**Figure S6.** Confusion matrix of the RF procedure on the validation set.

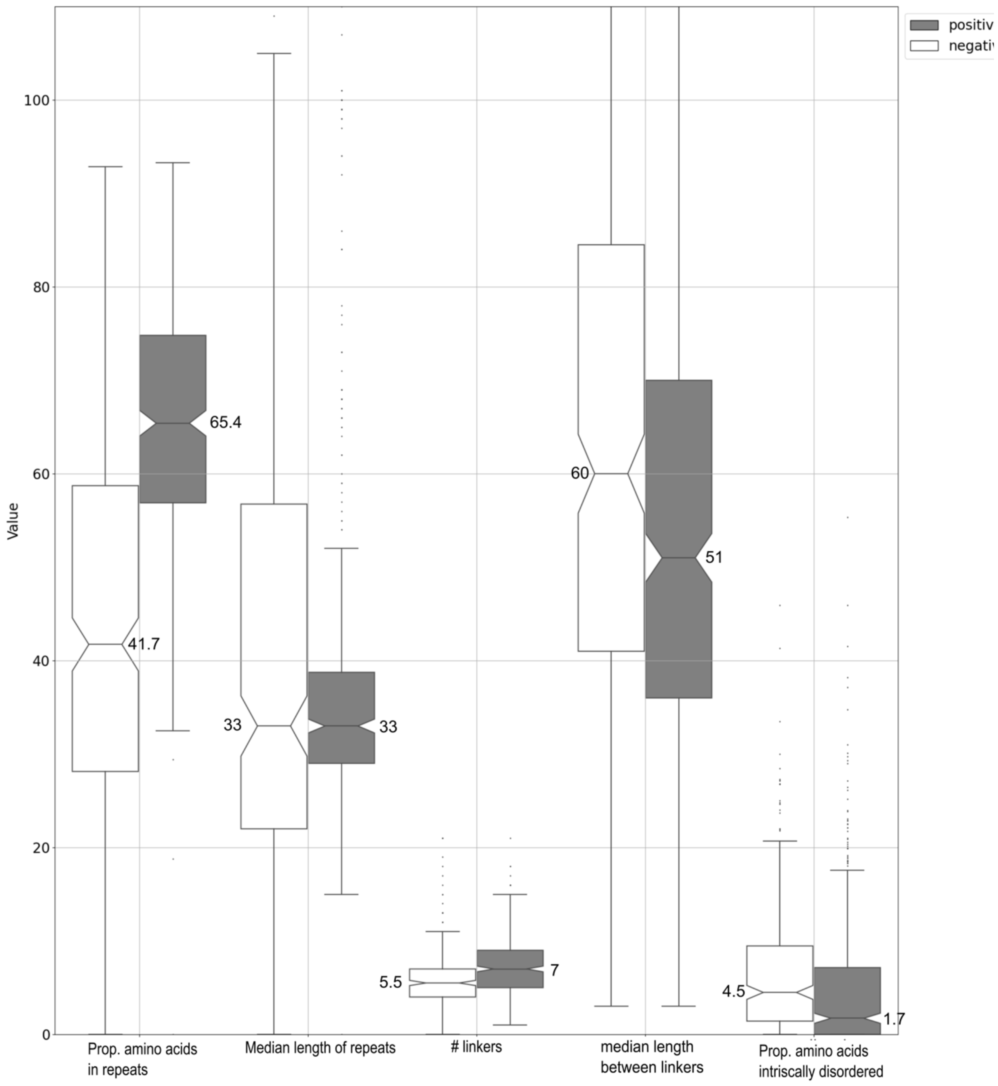


**Figure S6.** Barplot of the properties used in the DT and RF procedures within the positive (grey) and negative (white) proteins used in the training set. The median value is indicated for each distribution. All distributions are significantly different (chi-squared test, p-value < 0.05) between positive and negative controls, except for the median length of the repetition. The proportion of disorder is less in positive (0.02) than in negative control (0.04).
