## Supplementary material for "Searching α−solenoid proteins involved in organellar gene expression": Cattelin_TableSupp.pdf

| OTAF | Locus Id | Gene | Type | #motifs | Source |
|---|---|---|---|---|---|
| OPR, *Chlamydomonas reinhardtii* | Cre15.g638950 | NCC1 | NCL | 9 | (Boulouis et al. 2015) |
| | Cre15.g640400 | NCC2 | NCL | 9 | (Boulouis et al. 2015) |
| | Cre07.g348800 | TAB1 | Translation factor | 10 | (Rahire et al. 2012) |
| | Cre08.g358350 | TDA1 | Translation factor | 8 | (Eberhard et al. 2011) |
| | Cre06.g262650 | TAA1 | Translation factor | 7 | (Lefebvre-Legendre et al. 2015) |
| | Cre09.g394150 | RAA1 | Splicing factor | 4 | (Eberhard et al. 2011) |
| | Cre09.g388356 | TBC2 | Translation factor | 10 | (Eberhard et al. 2011) |
| | Cre06.g272450 | MBI1 | Maturation factor | 17 | (Wang et al. 2015) |
| | Cre10.g429400 | MCG1 | Maturation factor | 12 | (Wang et al. 2015) |
| | Cre16.g680850 | MDA1 | Maturation factor | 13 | (Viola et al. 2019) |
| | Cre10.g440000 | RAA8 | Splicing factor | 6 | (Marx et al. 2015) |
| | Cre09.g388372 | RAT2 | Maturation factor | 2 | (Marx et al. 2015) |
| PPR, *Arabidopsis thaliana* | AT2G18940 | - | P-class | 18 | (Pfalz et al. 2009) |
| | AT2G35130 | - | P-class | 12 | (Small and Peeters 2000) |
| | AT2G41720 | EMB2654 | P-class | 19 | (Lee et al. 2019) |
| | AT3G06430 | EMB2750 | P-class | 11 | (Williams and Barkan 2003) |
| | AT3G09650 | HCF152 | P-class | 12 | (Meierhoff et al. 2003) |
| | AT3G53170 | - | P-class | 10 | PPR database |
| | AT3G59040 | - | P-class | 10 | PPR database |
| | AT4G31850 | PGR3 | P-class | 27 | (Yamazaki et al. 2004) |
| | AT4G39620 | PPR5,EMB2453 | P-class | 9 | (Beick et al. 2008) |
| | AT5G02860 | - | P-class | 18 | PPR database |
| | AT5G48730 | - | P-class | 9 | (Williams and Barkan 2003) |

**Table S2.** OPR and PPR locus from which motifs were used to build the initial profiles for the IPB procedure, and associated references. Type refers of the OTAF function. P-class PPR contains an organelle targeting peptide followed by PPR motifs of 35 residues. They are involved in stabilization, processing, splicing and translation; PPR database: https://ppr.plantenergy.uwa.edu.au/.

| locus ID | mito/chlo | repeat radar | opr motif | α-solenoid shape | Annotation Phyto v5.6 | Annotation Phyto v6.1 |
|---|---|---|---|---|---|---|
| Cre01.g025808 | 2 | 12 | 1 | yes | - | OctotricoPeptide Repeat protein 122 |
| Cre01.g033250 | 2 | 8 | 1 | yes | - | - |
| Cre02.g109200 | 3 | 5 | 1 | yes | - | OctotricoPeptide Repeat protein 123 |
| Cre03.g145867 | 3 | 11 | 1 | yes, 7ptk | - | mitochondrial ribosome protein ml113 |
| Cre03.g168400 | 0 | 10 | 3 | yes* | - | OctotricoPeptide Repeat protein 134 |
| Cre03.g201103 | 2 | 9 | 1 | yes* | RAA7, psaA mRNA trans-splicing factor | psaA mRNA trans-splicing factor |
| Cre04.g217909 | 2 | 13 | 3 | yes | - | OctotricoPeptide Repeat protein 124 |
| Cre04.g228550 | 1 | 40 | 1 | yes * | - | - |
| Cre06.g251400 | 3 | 3 | 1 | no | MME6, NADP-dependent malic enzyme | NADP-dependent malic enzyme |
| Cre09.g395880 | 2 | 11 | 2 | yes | - | OctotricoPeptide Repeat protein 11 |
| Cre13.g579100 | 0 | na | 1 | yes * | APC1, Anaphase promoting complex subunit 1 | APC1, Anaphase promoting complex subunit 1 |
| Cre13.g605850 | 2 | 33 | 1 | yes * | - | OctotricoPeptide Repeat protein 127 |
| Cre13.g605900 | 2 | 28 | 1 | yes | - | OctotricoPeptide Repeat protein 128 |
| Cre16.g651900 | 2 | 12 | 1 | yes | CRI1, carotenoid isomerase | OctotricoPeptide Repeat protein 129 |
| Cre16.g662951 | 0 | 35 | 1 | yes | - | - |

**Table S3.1** Candidates retrieved by IPB unpublished and unreferenced as OPR with at least 2 motifs (OPR or $\text{repeat}_{\text{radar}}$). locus ID: locus identifier in the v5.6b annotation. targ: number of predictions for chloroplast/mitochondrial targeting; radar repeat: number of repeats predicted by RADAR. OPR motif: number of OPR motif found by IPB. α-solenoid shape: Yes/No (AF2 predictions). When an experimental 3D structure is available the PDB accession number is given. * indicates average pLDDT quality scores (< 0.5) for the AF2 predictions.

| Cluster id | # DTRF paralogs | Group | Species | InterPro annotation |
|---|---|---|---|---|
| 63 | 9 | Glaucocystophyceae | *Cyanophora paradoxa* | na |
| 99 | 7 | Rhodophyta | *Porphyridium purpureum* | IPR000477/Reverse transcriptase domain |
| 96 | 7 | Streptophyta | *Arabidopsis thaliana* | IPR005630/Terpene synthase, metal-binding domain |
| 51 | 10 | Chlorophyta | *Edaphochlamys debaryana* | na |
| 178 | 5 | Chlorophyta | *Edaphochlamys debaryana* | IPR002893/Zinc finger, MYND-type |
| 179 | 5 | Chlorophyta | *Edaphochlamys debaryana* | na |
| 180 | 5 | Chlorophyta | *Edaphochlamys debaryana* | IPR002893/Zinc finger, MYND-type |
| 50 | 10 | Chlorophyta | *Tetrabaena socialis* | IPR002893/Zinc finger, MYND-type |
| 175 | 5 | Chlorophyta | *Tetrabaena socialis* | IPR002893/Zinc finger, MYND-type |
| 176 | 5 | Chlorophyta | *Tetrabaena socialis* | IPR002893/Zinc finger, MYND-type |
| 177 | 5 | Chlorophyta | *Gonium pectorale* | IPR002893/Zinc finger, MYND-type |

**Table S8.** Species-specific protein families
